# Identification and Validation of Novel Microtubule Suppressors with an Imidazopyridine Scaffold through Structure-Based Virtual Screening and Docking

**DOI:** 10.1101/2021.12.08.471724

**Authors:** Samia A. Elseginy, A. Sofia F. Oliveira, Deborah K. Shoemark, Richard B. Sessions

**Author notes:** Corresponding Author: Richard B. Sessions, School of Biochemistry, University of Bristol, Biomedical Sciences Building, Bristol, BS8 1TD. UK.

## Abstract

Targeting the colchicine binding site of α/β tubulin microtubules can lead to suppression of microtubule dynamics, cell cycle arrest and apoptosis. Therefore, the development of microtubule (MT) inhibitors is considered a promising route to anticancer agents. Our approach to identify novel scaffolds as MT inhibitors depends on a 3D-structure-based pharmacophore approach and docking using three programs MOE, Autodock and BUDE (Bristol University Docking Engine) to screen a library of virtual compounds. From this work we identified the compound 7-(3-Hydroxy-4-methoxy-phenyl)-3-(3-trifluoromethyl-phenyl)-6,7-dihydro-3H-imidazo[4,5-b] pyridin-5-ol (**6**) as a novel inhibitor scaffold. This compound inhibited several types of cancer cell proliferation at low micromolar concentrations with low toxicity. Compound **6** caused cell cycle arrest in the G2/M phase and blocked tubulin polymerization at low micromolar concentration (IC_50_ = 6.1 ±0.1 μM), inducing apoptosis via activation of caspase 9, increasing the level of the pro-apoptotic protein Bax and decreasing the level of the anti-apoptotic protein Bcl2. In summary, our approach identified a lead compound with potential antimitotic and antiproliferative activity.

## 1. Introduction

Microtubules (MTs) consist of α/β tubulin heterodimers^1^ and are ubiquitous in all eukaryotic cells being the key component of the cytoskeleton.^2^ They play a critical role in many cellular processes, including cell division in which they assemble to make up the mitotic spindle required for segregation of the chromosomes to the spindle poles and consequent daughter cells; cell proliferation, maintenance of cell shape and signal transduction; and MT-motor proteins that transport diverse cellular cargoes.^3^ MTs are characterized by their highly dynamic behaviour, as they switch between periods of elongation and shortening.^4^ Targeting the process of microtubule dynamics is an excellent strategy for chemotherapy and modulation of MT dynamics is considered to be one of the most successful approaches in the treatment of cancer.^5^ Microtubule-targeting agents are classified into microtubule destabilizers and microtubule stabilizers according to the mechanism by which they affect microtubule dynamics.^6^ There are four major ligand binding sites identified on the microtubule namely: the vinca and colchicine sites where ligand binding induces microtubule destabilization and the taxane and peloruside /laulimalide sites where binding typically induces microtubule stabilization.^7^ Taxanes, vinca alkaloids, and colchicine all showed potent inhibition of cancer cell lines. However, colchicine showed limitations as an antitumor agent in clinical trials due to its narrow therapeutic window.^8^ While vinca alkaloids and taxanes are effective, they are also complex natural products that are difficult to synthesize and generally show poor bioavailability.^9-10^ In addition, the emergence of resistance to these drugs has been reported.^11-12^ Research has focussed on developing novel colchicine site inhibitors (CSI), since the molecular structure of known colchicine-site inhibitors is less complex than that of taxanes and vinca alkaloids.^13^ However, microtubule-destabilizing agents that bind at the colchicine-binding site and reach clinical trials had significant side effects, for example ZD6126 is a phosphate prodrug of N-acetylcolchinol (NAC) releasing the drug after administration and *in-vivo* hydrolysis. NAC binds to the colchicine binding site and inhibits tubulin polymerization, reducing the proliferating immature endothelial cells that line the tumour blood vessels and consequently inducing tumour cell death. ZD6126 reached phase II trials for metastatic renal cell carcinoma and induced necrosis in the tumours causing a large reduction in tumour cell yield after a single dose of ZD6126 but due to its cardiotoxicity, was withdrawn.^14^ Similarly, ABT-751 showed antitumor activities against a broad spectrum of cancers including those resistant to conventional chemotherapies. ABT-751 is an anti-tubulin agent with anti-vascular properties that is responsible for the dysfunction of tumour blood vessels. Despite administration of a single dose of ABT-751 (30 mg/kg, intravenously) disrupting tumour neovascularisation, it was withdrawn from Phase II due to adverse side effects.^15^

Despite the great potential of combretastatin and its prodrugs, CA1P and CA4P, these also suffer from drawbacks.^16^ As such, there remains an urgent need to design and discover novel MTs inhibitors based on the colchicine site of β-tubulin.^17^ Therefore, our objective is to identify a novel chemical scaffold to bind the colchicine site that may form the basis of a new lead compound which offers promising antimitotic and antiproliferative activity as well as is well-tolerated. Here we have used structure-based pharmacophore virtual screening of a subset of the ZINC15 database.^18^ This computational approach represents a quick and efficient method in the identification of novel and diverse scaffolds of CSI. Molecular docking was carried out with three programs^19^; the Bristol University Docking Engine (BUDE) ^20,21^, AutoDock 4.2 ^22^ and MOE (https://www.chemcomp.com/). Selected hits from this procedure were assessed for antiproliferative activity via their ability to cause mitotic spindle arrest, affect tubulin polymerization and induce apoptosis.

## Results and Discussion

### Identifying colchicine binding site inhibitors

The binding orientation of colchicine was unequivocally established in the structural determination of the tubulin/DAMA-colchicine complex due to the electron density of the sulphur atom (PDB: 1SA0).^23^ Hence, a 3D pharmacophore was built based on the analysis of the interaction of colchicine with tubulin in this structure. The derived pharmacophore (Figure S1) consists of seven features^24^ comprising three H-bond acceptor centres corresponding to the interactions with Cysβ241 and Val α181, one H-bond donor, an aromatic centre, and two hydrophobic centres corresponding to interactions with Leuβ248, Alaβ250, Leuβ255, Asnβ258, Alaβ316, and Valβl318 (Zone 2).^25^ In order to explore this approach we chose the first one hundred thousand compounds from the current 9.9 million clean, drug-like and purchasable compounds in the ZINC15 database and filtered this set against the 3D pharmacophore using MOE. This process afforded 2476 compounds matching at least 4 points of the pharmacophore. This set of compounds and the native ligand (colchicine) were docked with BUDE, MOE, and AutoDock 4.2 into the colchicine binding site. MOE and Autodock4.2 are used widely in computational studies, BUDE is an in-house docking program using an empirical free energy forcefield to predict ligand affinities and has been used for inhibitor discovery^26-29^, The predicted binding affinities of colchicine using BUDE, MOE, and AutoDock4.2 (−100.27 kJ/mol, -5.1 kcal/mol and -9.52 kcal/mol) respectively, were used as a cut-off for selecting potential hits. The numbers of compounds passing this filter with a binding score better than colchicine were 188, 107, and 226 respectively. Next, we applied the criterion that compounds must be common to two or three of these docked sets. A total of 99 compounds passed this selection step and were assessed for toxicity risk using Osiris Property Explorer (https://openmolecules.org), resulting in the exclusion of a further 38 compounds (ESI Properties_99-compounds.xlsx). The remaining 61 compounds were clustered using the Flexophore descriptor implemented in the DataWarrior cheminformatics application (Figure S2) (https://openmolecules.org). A shortlist was generated by sampling from the compound clusters and the final shortlist of 13 compounds was made using actual compound availability and using cost as a proxy for synthetic accessibility. Of the thirteen shortlisted, only compound **3** is reported in the scientific literature^30^ according to a search using the Chemical Abstracts service, SciFinder. The similarity of the compound set with the 50 CSI ligands currently present as tubulin complexes^31^ in the PDB was assessed using DataWarrior and the results (Figure S3) with the Flexophore descriptor^32^ show partial overlap at the 60% level. These 13 compounds all passed the PAINS filters (http://zinc15.docking.org/patterns/home and http://www.cbligand.org/PAINS) and were purchased from MCULE (https://mcule.com) for experimental evaluation (Figure 1, Tables S1 and S2).

**Figure 1.**
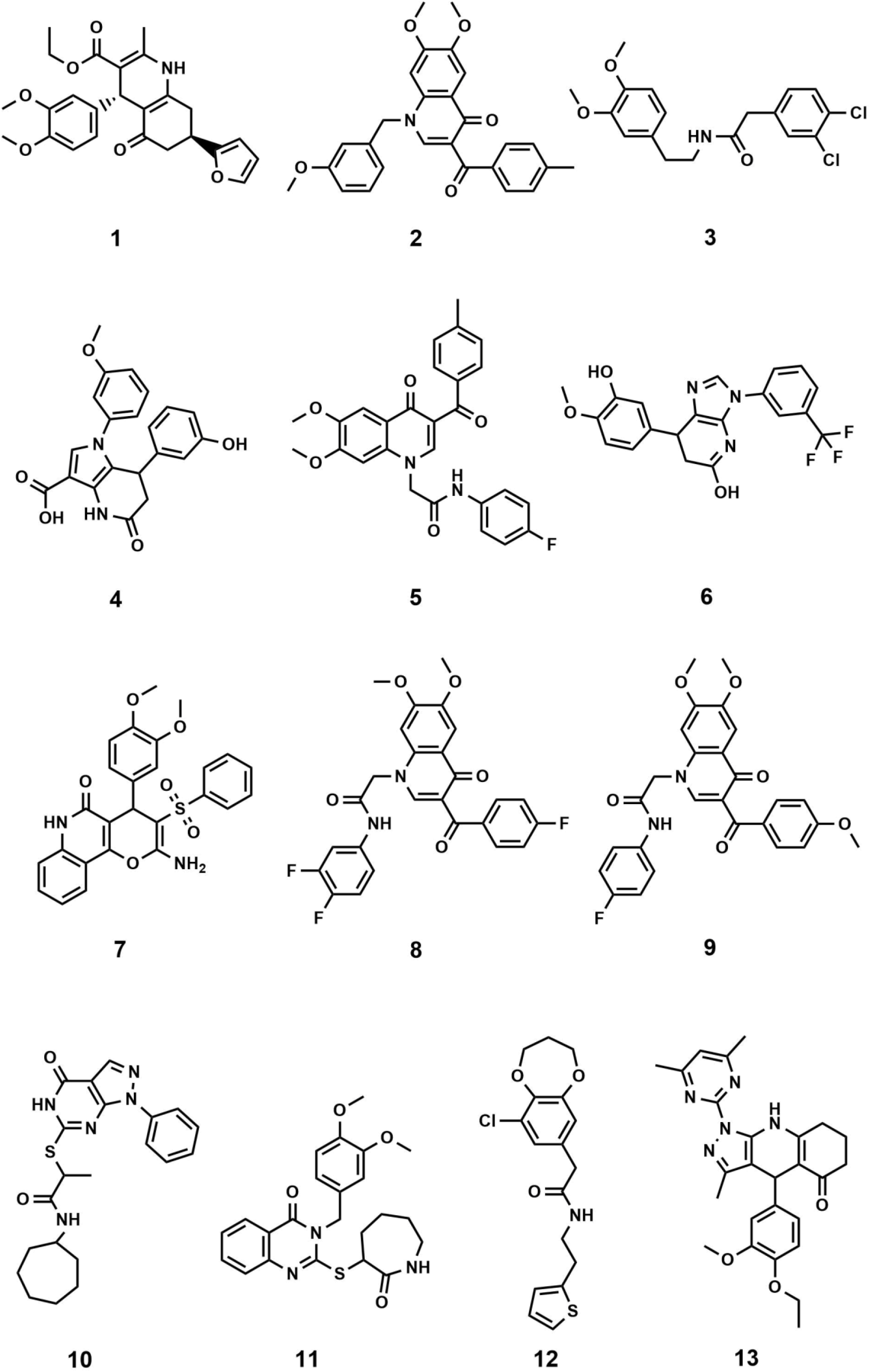
Structures of the 13 selected compounds

### Structure characteristics of the shortlist compounds

In all but one case the selected compounds are characterised by having core structures comprising two or three fused cyclic rings with either some or extensive aromatic character (Figure 1). The exception is compound **3** which consists of two substituted phenyl rings coupled by a five-atom long linker. The core structures are fused bicyclic rings: [5,6,0] (**4**,**6**,**10**), [6,6,0] (**1**,**2**,**5**,**8**,**9**,**11**); [6,7,0] (**12**) and fused tricyclic structures with ring sizes 6:6:6 (**7**) and 5:6:6 (**13**) and all are N or O heterocyclic systems including quinolines and quinolones. The core structures are typically elaborated with two phenyl groups, either attached directly or via short linkers to give compounds with moderate flexibility. Four compounds (**10**,**11**,**12**,**13**) have heterocyclic rings attached to their cores. All but two compounds (**10**,**12**) have at least one methoxy-aryl substituent, four (**5**,**6**,**8**,**9**) are fluorinated and one (**3**) has a dichlorophenyl group.

## Biological Evaluation

### Antiproliferative activity

The anti-proliferation activities of the shortlisted 13 compounds were evaluated against MCF-7, MDA-231, and A549 cancer cells lines using the MTT assay^33^ at 10 μM in comparison to paclitaxel (16 nM), employed as a positive control. The results are shown in Figure 2A and indicate that **6, 8, 9, 13** showed significant anti-proliferative activities against all three cell lines. Compound **6** caused 50, 40, and 60% inhibition of MCF-7, A549, and MDA-231 cell growth, respectively. Similarly, **8** caused 50, 20, and 80% inhibition of MCF-7, A549, and MDA-231 cell growth, respectively. The best activity observed was against MDA-231, a triple-negative cell line that is highly aggressive and resistant to treatment. Compounds **9** and **13** caused 50 and 40% inhibition of all tested cancer cell types respectively, with limited toxicity to normal fibroblast F180 cells (Figure 2B).

**Figure 2.**
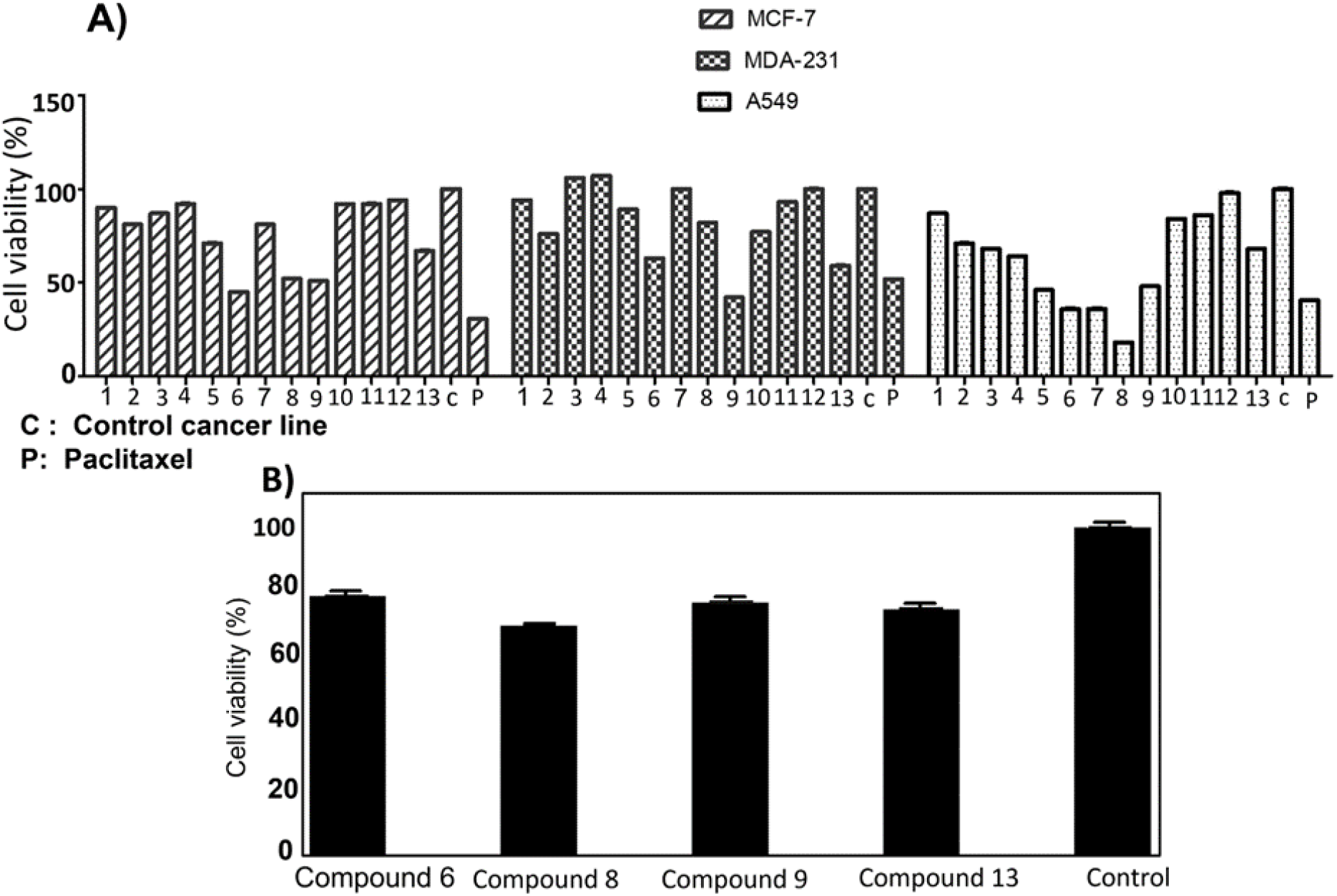
A) Antiproliferative activity of compounds **1 - 13** identified from the virtual screening against MCF-7, MDA-231 and A549 cell lines. B) Cytotoxic screening of compounds **6, 8, 9** and **13** against FibroblastF180 mammalian cell line showed that compound **6** with highest % viability which indicates its low toxicity. Control: Fibroblast F180 mammalian cell line.

The most active four compounds **6, 8, 9** and **13** were titrated against the three cell lines to determine their IC_50_ values in this cell assay (Figure 3). Compounds **6, 8**, and **9** showed IC_50_ values in the 9-20 μM range against MCF-7, A549, and MDA-231 while **13** showed IC_50_ values above 20 μM (Table 1) hence **6, 8** and **9** were chosen for more detailed investigation.

**Figure 3.**
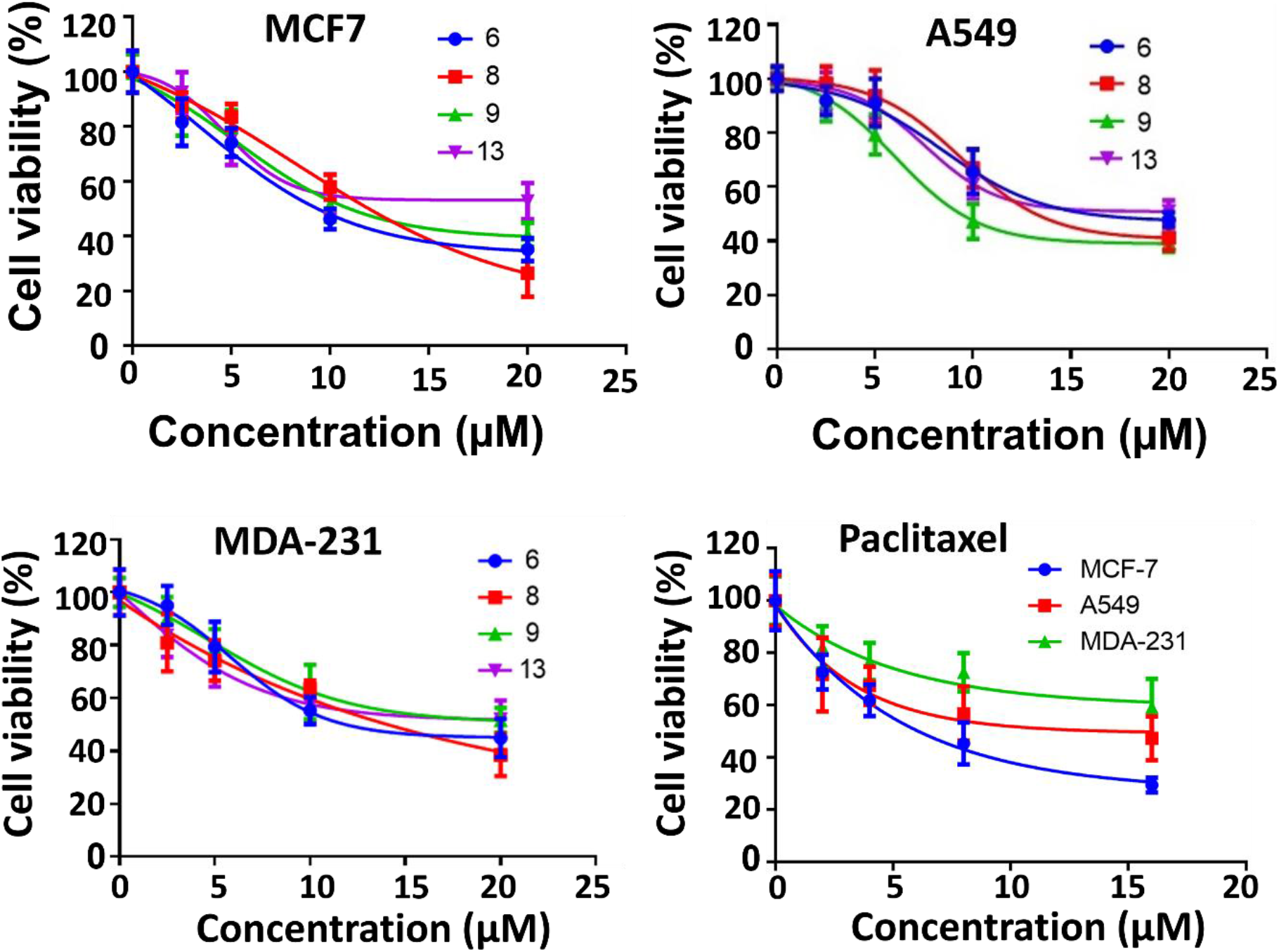
IC_50_ of the active compounds **6, 8, 9** and **13** against MCF-7, MDA-231 and A549 cancer cell lines, Paclitaxel was used as positive control. Each result is a mean of triplicate experiments, and the mean values and standard error of mean are shown.

**Table 1.** IC_50_ values of antiproliferative activity of compounds against MCF-7, A549, and MDA-231 cancer cell lines.

| Compound | MCF7 | A549 | MDA-231 |
| --- | --- | --- | --- |
| **6** | 9.4 ± 0.1 (μM) | 16.1 ± 0.3 (μM) | 11.9 ± 0.4 (μM) |
| **8** | 12.0 ± 0.5(μM) | 13.2 ± 1.1(μM) | 14.1 ± 1.0 (μM) |
| **9** | 11.0 ± 0.1 (μM) | 9.4 ± 0.6 (μM) | 19.5 ± 0.7 (μM) |
| **13** | 23.2 ± 2.4 (μM) | > 20 ± 4 (μM) | > 23 ± 3 (μM) |
| Paclitaxel | 6.1 ± 1.0 (nM) | 15.0 ± 5.0 (nM) | 19.3 ± 1.0 (nM) |

\*IC<sub>50</sub> values are the mean of three replicate experiments ± SD.

### Inhibition of Mitotic Spindle Formation

An immunofluorescence assay was used to investigate the mechanism of action of **6, 8**, and **9** on tubulin organization into mitotic spindles during cell division.^34^ All three compounds caused the formation of classical multipolar spindle profiles (Figure 4), similar to the positive controls paclitaxel and colchicine. These results indicated that compounds **6, 8**, and **9** caused modulation of tubulin assembly with irregular morphology, showing typical mitotic arrest.

**Figure 4.**
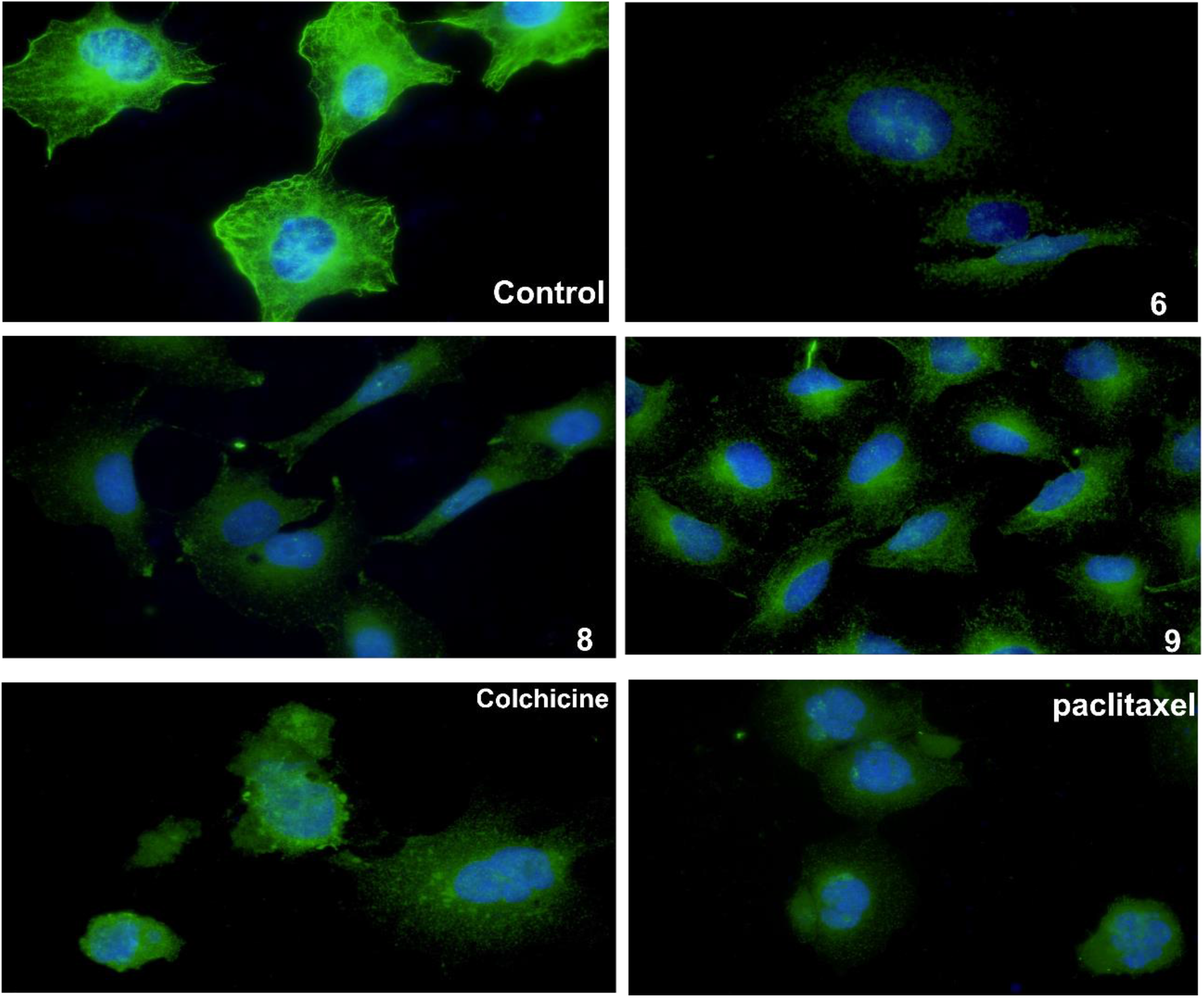
The three compounds **6**,**8** and **9** disrupt microtubule formation in A549 cells. A549 cells were treated for 24 h with DMSO as control, Paclitaxel (14 nM), Colchicine (0.1 nM), compound **6** (16μM), compound **8** (13μM), compound **9** (9 μM), then fixed, and stained with anti-β-tubulin antibody (green) and with DAPI for DNA (blue) to visualize the microtubules.

### Tubulin Polymerization Assay in vitro

Microtubule polymer solutions scatter light in a concentration-dependent manner.^35,36^ This behaviour was used to monitor the effect of ligands on microtubule polymerization (Figure 5). Compounds **6, 8**, and **9** at 15 μM and paclitaxel and colchicine at 3 μM were incubated with unpolymerized tubulin protein at 37 °C. The tubulin polymerization activities were determined by measuring the fluorescence and recording the area under the curve (AUC). Increasing fluorescence indicates increasing polymerization activity while decreasing the fluorescence indicates greater depolymerization activity. Paclitaxel which stabilizes polymerized tubulin caused an increase in the AUC by ∼1.3 fold. On the other hand, the destabilizing compound, colchicine, caused a decrease the AUC by ∼1.3 fold when compared to the negative control, hence inhibiting polymerization of tubulin. Compound **6** showed a modest decrease in fluorescence in comparison to the control while **9** increased the AUC by a similar amount and **8** showed an AUC similar to the control. Although these results are within error, they suggest that **6** inhibits and **9** stabilises tubulin polymerisation (Figure 5 A and B).

**Figure 5.**
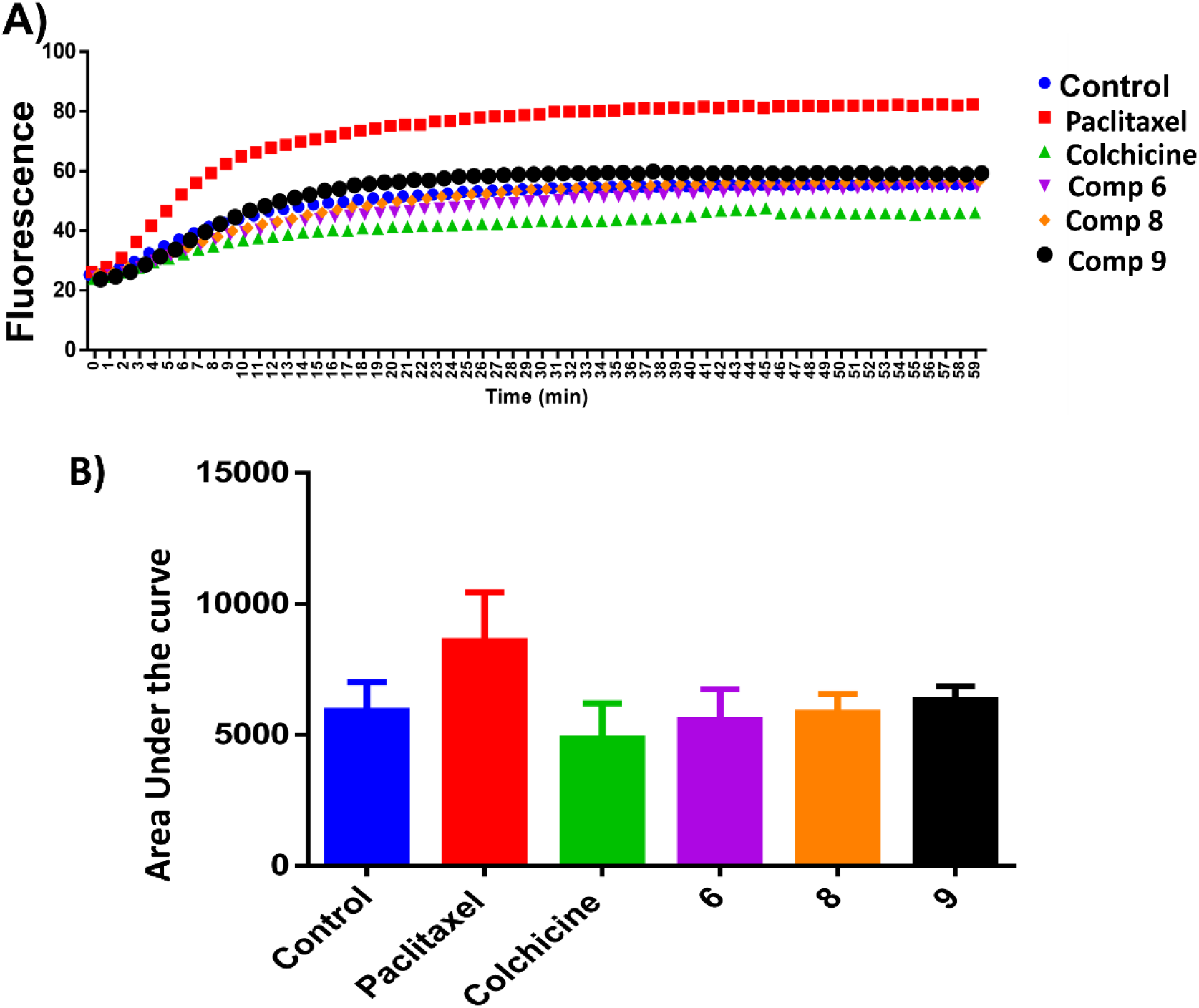
A) Tubulin polymerization of compounds **6, 8** and **9**. Tubulin polymerization was monitored by the increase in the fluorescence at 360 nm (excitation) and 420 nm (emission) for 1 h at 37 °C. Paclitaxel and colchicine were used as the positive control while 0.1 % DMSO used as negative control. B) Area under the curve for the tested compounds and positive and negative controls.

### Tubulin polymerization inhibition mechanism in cells

An ELISA assay was used to measure tubulin polymerization in MCF7 cells in the presence of compounds **6, 8**, and **9**. The results are shown in Figure 6 and indicate that **6** behaves like colchicine as a suppressor of microtubule polymerization, while **8** and **9** enhance polymerization. These results are in reasonable accordance with the in-*vitro* tubulin polymerization assay results and are more reliable, showing differences well outside the standard error. IC_50_ values of Compounds **6, 8**, and **9** required to modulate tubulin polymerization were 6.1 ± 0.1, 13.1 ± 0.3, and 12.8 ± 0.2 μM respectively (Figure 6, Table 2). Compound **6** was selected for further investigation due to its low toxicity on normal cells while affording good potency against cancer cells (particularly MCF7 cells). It also showed the best activity in the tubulin polymerization assays. Although the IC_50_ of compound **6** is a little higher than colchicine (6.1±0.1 μM and IC_50_ 1.4±0.02 μM respectively, Table 2) the low cytotoxicity of this compound suggests that it is a promising lead-molecule and scaffold for drug development.

**Figure 6.**
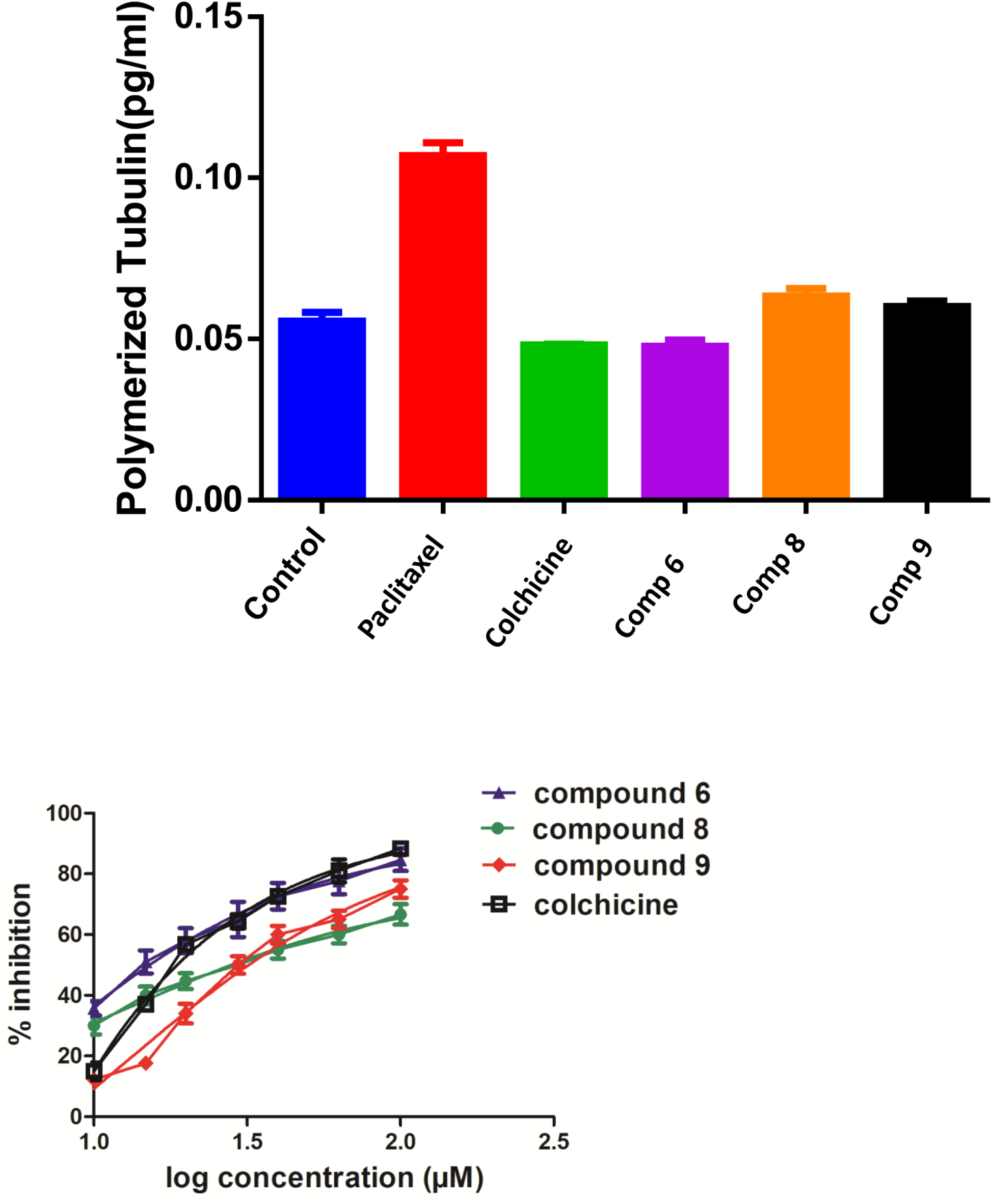
ELISA binding assay. A) MCF7 cell line treated with compounds **6**,**8** and **9** at their IC_50_ for 24h, paclitaxel and colchicine were used as positive controls. The tubulin polymer was extracted, and the quantities of monomeric and polymeric tubulin were measured using ELISA. Compound **6** showed the same effect as colchicine. B) IC_50_ of the active compounds **6, 8, 9** and colchicine in tubulin polymerization assay. Each result is a mean of triplicate experiments, and the mean values and standard error of mean are shown.

**Table 2.** IC_50_ value of compounds affecting tubulin polymerization in MCF-7 cells.

| Compounds | IC<sub>50</sub> (μM) |
| --- | --- |
| **6** | 6.1 ±0.1 |
| **8** | 13.1±0.3 |
| **9** | 12.8±0.2 |
| Colchicine | 1.4±0.02 |

\*IC<sub>50</sub> values are the mean of three replicate experiments ± SD.

### Compound 6 Inhibited Cell Cycle Progression at G2/M and induced apoptosis

Cell cycle analysis was performed to determine at which phase compound **6** exerted its antimitotic effect. MCF7 cells were treated with compound **6** at its IC_50_ concentration (6 μM) and 0.1% DMSO as control and incubated for 24 h, followed by measuring cell cycle distribution by flow cytometry. The results showed that the antimitotic activity of compounds **6** was through apoptosis. Compound **6** caused a 10-fold increase in cell populations at the G2/M compared to control (Figure 7A-C, Table S3). The apoptotic activity of compound **6** was further evaluated by propidium iodide (PI) and annexin-V-FITC labeling assay on MCF-7 cells and using flow cytometry analysis^37^. Compound **6** caused an 84-fold increase in the late stage of cellular apoptosis compared to the negative control (Figure 7D-F). Furthermore, the compound also caused a decrease in early apoptosis by 11-fold compared to the negative control (Figures 7D-F).

**Figure 7.**
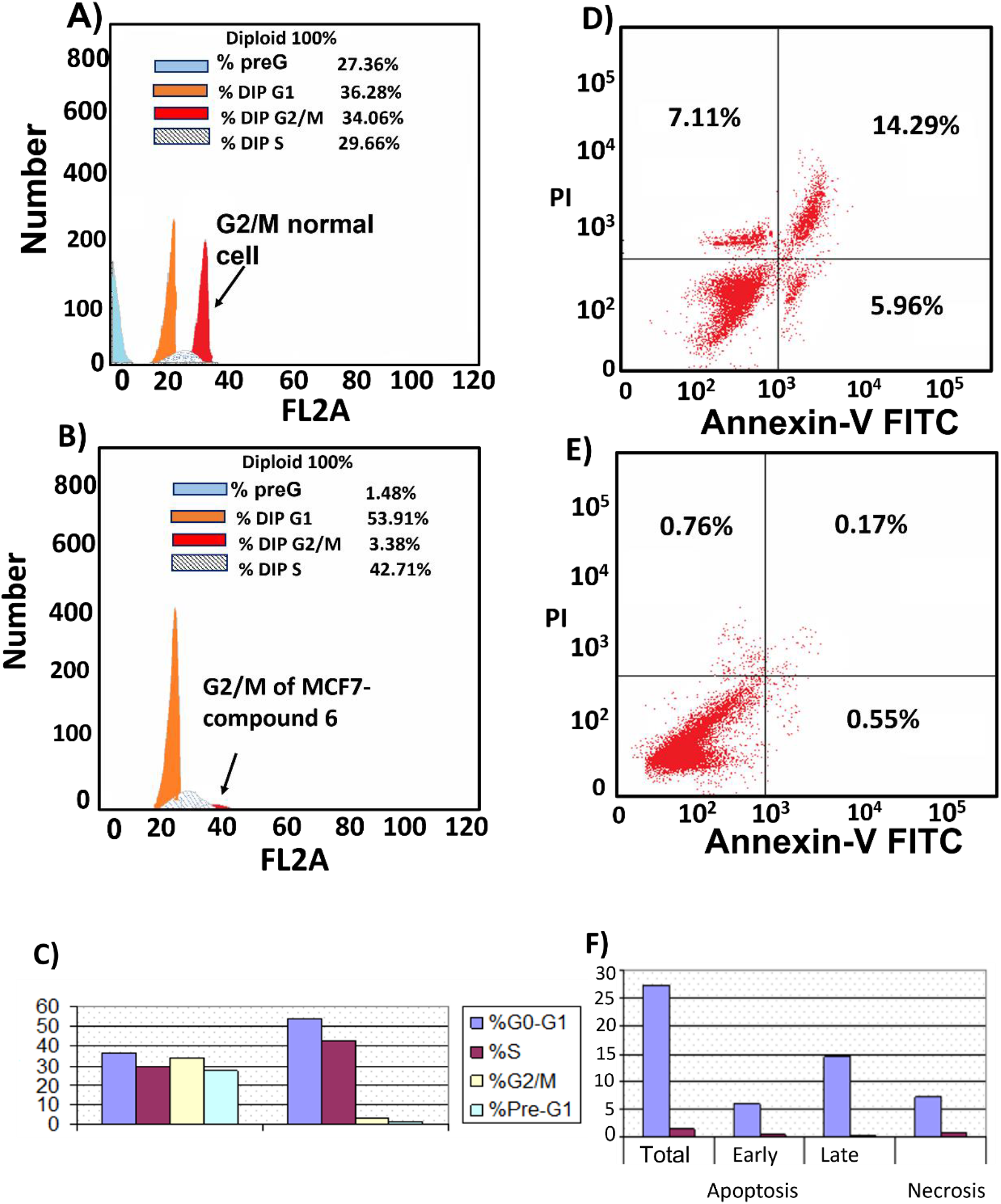
Cell cycle distribution. MCF-7 cells were treated with A) 0.1 % DMSO (B), compound **6** (6 μM) for 24 h. Next, the cells were harvested and stained with propidium iodide, and flow cytometry cell cycle analysis was used to evaluate the cell cycle progression. C) Percentages of cells in the different phases indicate compound **6** arrests the cell cycle at (G2/M). Cell apoptosis D) MCF7 treated with compound **6** for 24 h showed late apoptosis. E) MCF7 treated with 0.1 % DMSO as control. F) Percentage of MCF7 cell apoptosis showed that compound **6** increased both stages of apoptosis with a large increase in late apoptosis.

**Figure 9.**
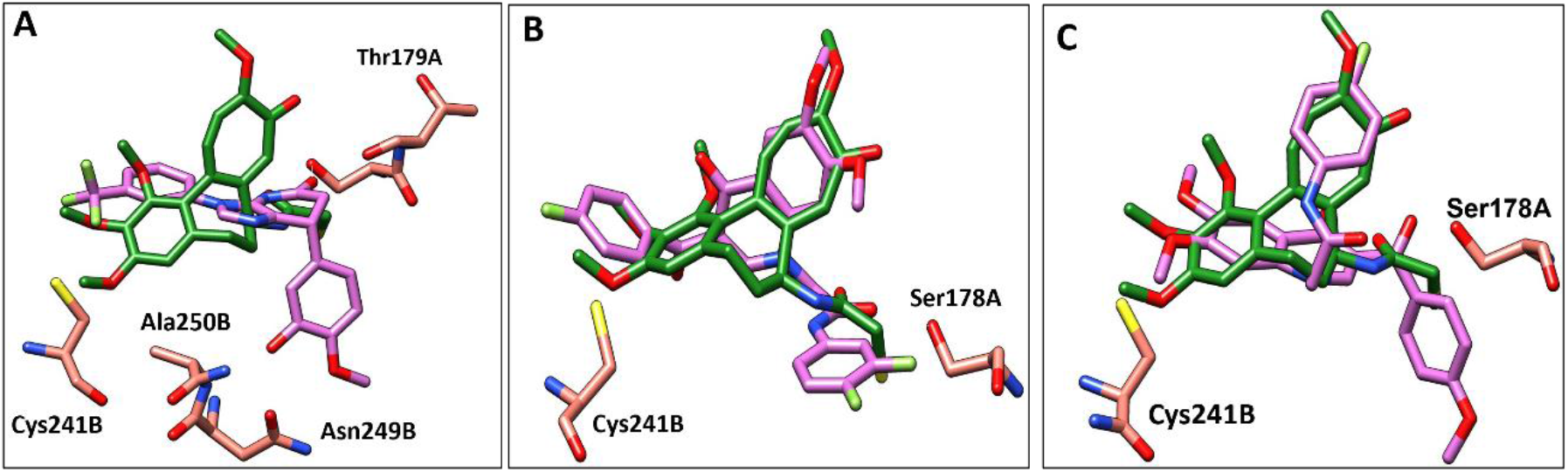
Comparison of the proposed binding modes of compounds 6, 8 and 9 into α/β interface of tubulin (PDB:1SA0) with DAMA-colchicine binding mode (dark green sticks). A) compound **6** (purple sticks), B) compound **8** (purple sticks) C) compound **9** (purple sticks).

**Figure 10.**
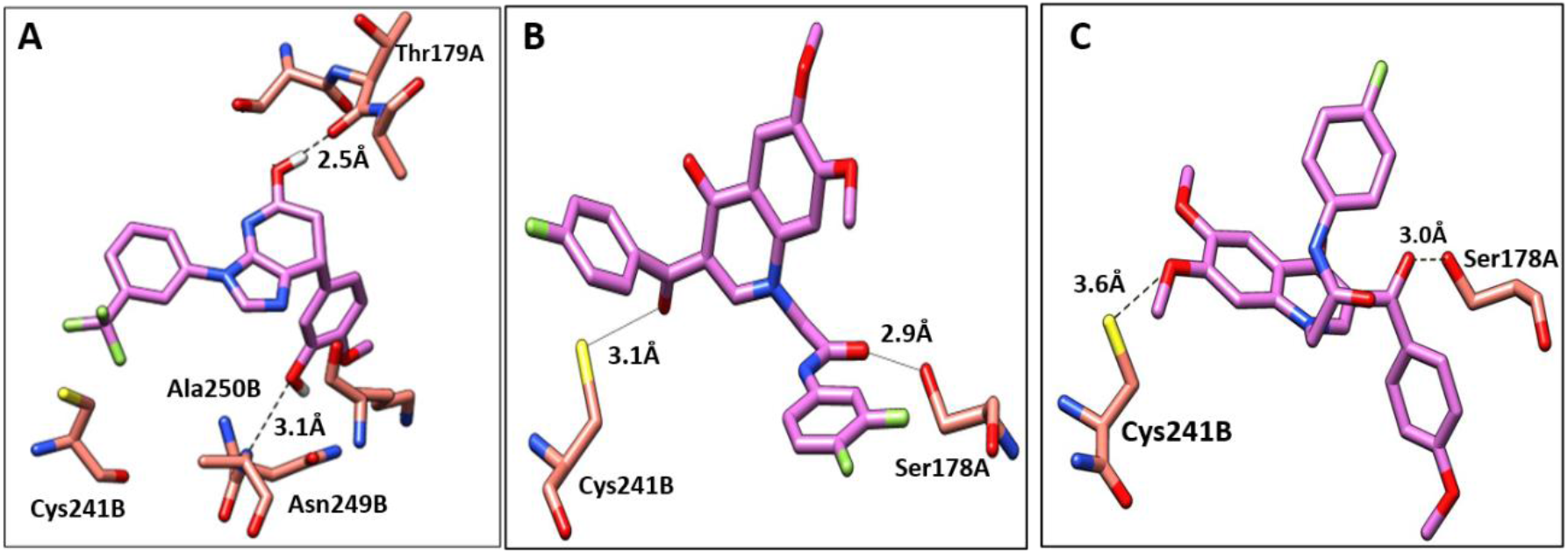
The interactions of compounds **6, 8**, and **9** with tubulin dimer (PDB:1SA0). A) Compound **6**, B) Compound **8**, C) Compound **9**, hydrogen bonds are shown as dotted lines.

### Compound 6 acts by a dual apoptosis mechanism

The apoptotic effect of compound **6** was determined by measuring caspase 9 levels. Caspase 9 initiates apoptosis by activating a cascade of intracellular events.^38,39^ MCF7 cancer cells were treated with compound **6** at 6 μM for 24 h. The level of caspase 9 was then measured by Enzyme-Linked Immuno-Sorbent assay (ELISA) analysis. Compound **6** showed an increase in the level of caspase-9 in treated cells by 7.6-fold compared to control-vehicle-treated cells (Table 3). The Bcl2 family of proteins plays a crucial role in regulating mitochondrial apoptosis by either increasing the level of Bcl2 anti-apoptotic proteins or downregulating the level of Bax, a pro-apoptotic protein. Hence, cancer cells may develop resistance to apoptosis by changing the level of the Bcl2 and Bax protein expression. The effect of compound **6** on the balance of Bcl2/Bax proteins in MCF7 cancer cells was investigated. The results showed that compound **6** increased the level of the pro-apoptotic protein Bax by 7.2-fold compared to the control, and at the same time decreased the level of the anti-apoptotic protein Bcl2 by 2.7-fold (Table 3).

**Table 3.** Determination of Caspase 9, Bcl2, and Bax levels in the MCF7 cells treated with compound **6**

| compounds | Caspase9 (ng/ml) | Bcl2 (ng/ml) | Bax (pg/ml) |
| --- | --- | --- | --- |
| **6** | 13.7±0.7 | 2.1±0.1 | 207.6±6.3 |
| control | 1.8±0.3 | 5.7±0.1 | 28.8±1.6 |

\* Values are the mean of three replicate experiments ± SD.

### Compound verification

^1^H NMR spectra of compound **6** (Figure 6), compounds **8** and **9** (Supplementary Figures S6-S11) were measured to confirm the chemical structures. Likewise, molecular weights were confirmed by mass spectroscopy for compound **6** MS m/z = 404.36 [M^+^+1] and compound **8** MS m/z

497.37 [M^+^+1] (Supplementary Figures 2,4 respectively).

## Computational Modelling

### Molecular Docking

To further elucidate the interactions between compounds **6, 8** and **9** and tubulin, the binding mode of the three compounds within the colchicine binding site was investigated. The results of docking **6, 8**, and **9** are summarized in Table S4. The docking study revealed that the compounds fitted well within the hydrophobic pocket of β-tubulin, making hydrophobic interactions with the hydrophobic residues lining zone 2 of the colchicine binding pocket, namely Leuβ242, Leuβ248, Alaβ250, Leuβ255, Valβl315, Alaβ316, and Ileβ378. The trifluoro phenyl ring of compound **6** occupied the same position as the trimethoxy phenyl ring of colchicine, facilitating hydrophobic interactions with key residues (Figure 7A). In the case of compound **8**, the 4-fluoro phenyl ring occupied the position of the trimethoxy phenyl ring of colchicine and the dimethoxy quinoline ring adopted an analogous position as the tropone ring of colchicine (Figure 7B). However, compound **9** showed a different binding mode with the dimethoxy quinoline ring positioned where the trimethoxy phenyl ring of colchicine and the 4-fluoro phenyl group of tropone (6C) reside. The three compounds showed strong H-bonds, as the OH of compound **6** served as H donor and formed an H-bond with Thrα179 and OCH_3_ formed an H-bond with NH-Alaβ250 (Figure 8, and Table S4). Both compounds **8** and **9** formed H-bonds with Serα178, and Cysβ241. (Figure 8, and Table S4). The trifluoro phenyl group of compound **6** faced SH-Cysβ241, allowing the lone pair of electrons of the sulphur to interact with the pi cloud of the aromatic ring (SH---π).^40^ Analyses of the docking results indicate that the hydrophobic interactions of compounds **6, 8** and **9** predominate while H-bonds help in the proper orientation of the compounds within the binding pocket.

### Molecular dynamics

Simulations of the tubulin-ligand complexes (colchicine, **6, 8, 9, 14** and **15**) were performed for 3 repeats of 500 ns with different initial starting velocities to assess their stability and persistence over this short period. The Root Mean Square Deviation (RMSD) of the protein and the ligand with respect to their initial (time 0 ns) positions are shown in Figure S4 and average values in Table S5. In all cases the RMSD of the protein stays between 0.2 and 0.3 nm for 99.15 % the 9 us of total simulation time (maximum 0.34 nm). Colchicine and **6** also stay close to their original positions in all simulations with average RMD values of 0.15 nm (maximum 0.32 nm) and 0.14 nm (maximum 0.28 nm) respectively, indicating that the conformation and poses of these complexes are close to their original docked positions. The behaviour of **8** is more labile with the ligand RMSD values typically ranging between 0.3 and 0.4 nm. In these simulations, the 4-fluorophenyl group reorients closer to the deep-binding site (by Y224 and V238) the naphthyl group tilts by some 45° in two simulations while remaining close to the original pose in run 3. The greatest variability in pose and conformation occurs in the 3,4-difluorophenyl group at the dimer interface. Compound **9** stays close to its original docked position in runs 1 and 3 while in run 2 it rapidly slips the 4-methoxyphenyl ring into the deep-binding pocket causing the whole ligand to shift by some 0.3 nm into the B subunit, accounting for the large but stable rise in RMSD to 0.4-05 nm in this case. Compounds **14** and **15** are versions of **6** redesigned to occupy the deep-binding pocket (see below) and their plots of RMSD versus time indicate little movement from the original pose as seen for colchicine and **6**. Ligand-residue contact data are provided for the 500 ns structures from the 12 simulations of tubulin with **6, 8, 9** and colchicine in Figure S5.

### Rational design and synthetic accessibility

Many of the protein ligand interactions between **6** and tubulin are mediated by the two aryl groups attached to the imidazopyridine core. A simple synthetic route to varying these substituents has been described^41^ and is shown here.

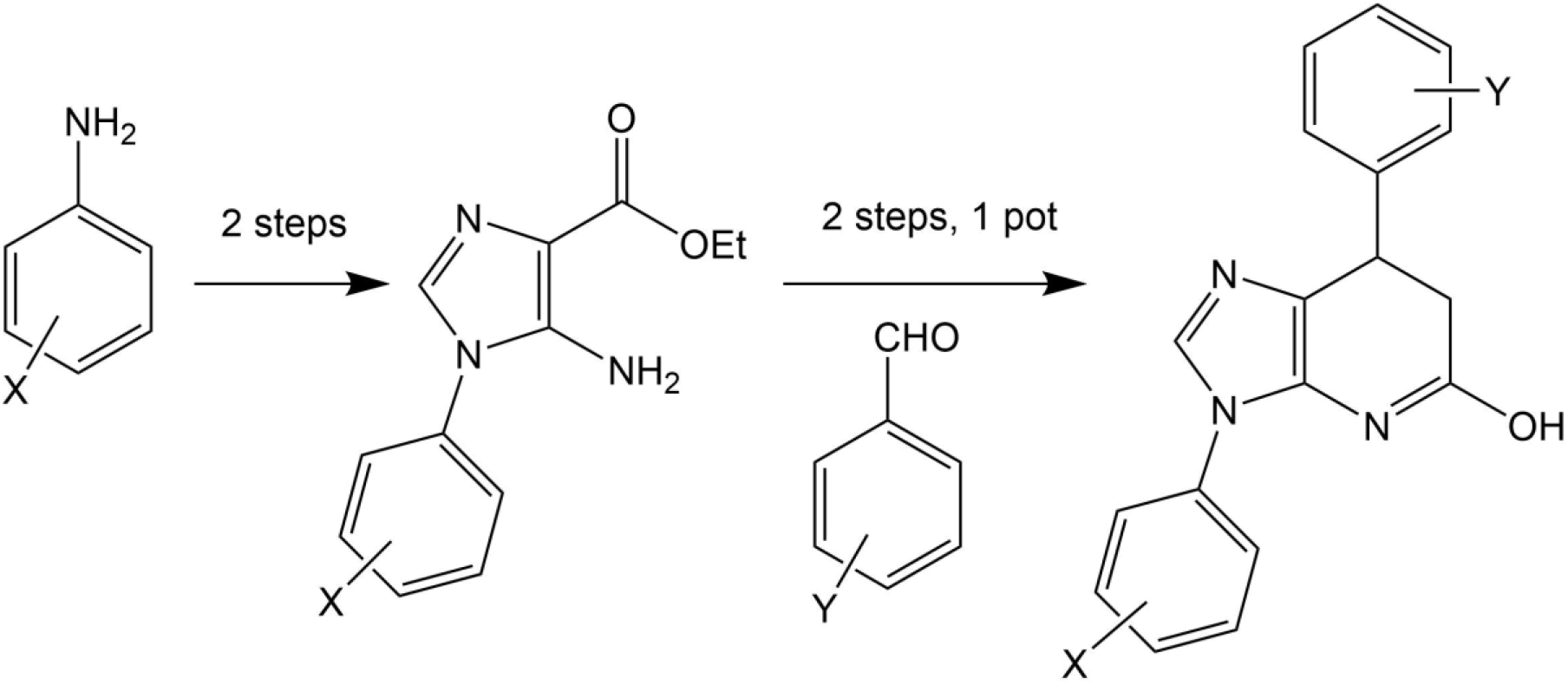

The mild condition of this route and wide variety of substituted aniline and benzaldehyde starting materials available commercially make this a very attractive goal for medicinal chemistry to explore SAR and rational design for improving affinity and the drug-likeness of this potential lead. As an exemplar, we describe the design and modelling of **14** with variations to exploit the deep-binding pocket (X_1_ = 3-hydroxy; X_2_ = 4-pyrimidine) and improve interactions at the dimer interface (Y = 3,5- bistrifluoromethylbenzene).

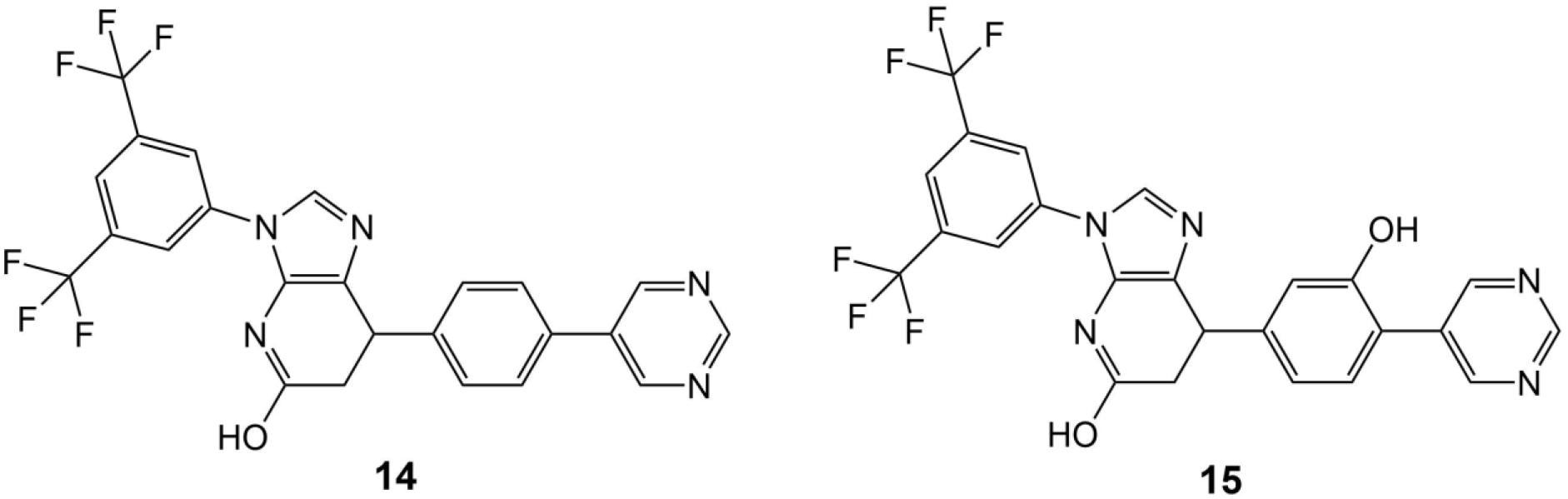

We also investigated **15**, equivalent to **14** with X_1_ = 3-hydroxy removed, since this is cheaper to synthesise and in conjunction with **14** would allow exploration of any contribution of the binding affinity of this group that hydrogen bonds with the carbonyl oxygen of V238 in our modelling.

## Conclusion

In this paper we have used computational methods to identify inhibitors and modulators of tubulin polymerization. Firstly, 7-feature pharmacophore matching was used to filter out some 98 % of compounds from a library of 100,000 available compounds. Consensus docking of these 2746 matched compounds was performed using three different methods (MOE, BUDE and Autodock) and passed 99 compounds. Cheminformatics was used to remove compounds with toxicological risk and to cluster compounds by similarity. A total of 13 compounds were chosen from the clustering for experimental investigation. The antiproliferative activities of the 13 compounds were evaluated against three cancer cell lines (MCF-7, MDA-231, and A549) and four compounds (**6, 8, 9** and **13**) showed significant results. Compounds **6**,**8** and **9** had IC_50_ values ≤ 20 μM and these three compounds disrupted spindles in the mitotic cells giving a phenotype similar to colchicine. Compounds **6**,**8** and **9** modulated tubulin polymerisation in vitro and in three cancer cell lines with minimal toxicity. Molecular dynamics simulations into the microsecond regime support the predicted binding modes. Compound **6** showed the lowest IC_50_ (6.1±0.1 μM) inhibiting tubulin polymerization in MCF-7 cells using ELISA. FACS cell cycle analysis showed that **6** arrests the cell cycle at the G2/M phase and induces late apoptosis via upregulating caspase-9 and Bax while downregulating Bcl2. The activity of **6** against cancer cells, its low cytotoxicity to normal fibroblasts and ease of synthesis of variants provides a start-point for medicinal chemistry development. The results of docking and simulation of **6** were used to suggest elaborations to exploit the deep-binding site in the colchicine pocket and the persistence of binding of these derivatives (**14** and **15**) was demonstrated by further molecular dynamics simulations. The proposed synthetic route should facilitate production of a large number of derivatives based on the imidazopyridine scaffold of **6** to explore SAR and improve the drug-likeness of this series.

## Experimental Section

### Chemistry

^1^H NMR spectra were recorded on a Bruker spectrometer at 500 MHz. Chemical shifts are expressed in parts per million (ppm) relative to tetramethylsilane and coupling constants (J values) are represented in Hertz (Hz) and the signals are designated as follows: s, singlet; d, doublet; t, triplet; m, multiple. Mass spectroscopic data were obtained through Electrospray ionization (ESI) mass spectrometry.

7-(3-Hydroxy-4-methoxy-phenyl)-3-(3-trifluoromethyl-phenyl)-6,7-dihydro-3H-imidazo[4,5-b] pyridin-5-ol (**Compound 6)**. ^1^H NMR (DMSO-d6) δ: 2.72 (m, 1H, <u>H</u>-CH), 3.12 (m, 1H, H-C-<u>H</u>), 3.75 (s, 3H, OCH_3_), 4.23 (m, 1H, CH), 6.25 (m, 2H, ArH), 6.80 (d, 1H, J= 8, ArH), 7.54 (d, 1H, J= 6.5, ArH), 7.82 (m, 4H, ArH, imidazole C<u>H</u>), 8.60 (s, 1H, OH), 10.2 (s, 1H, OH). ^13^C NMR (DMSO-*d6*): δ 35.59 ppm (CH of the pyridine ring), δ 38.44 ppm (CH_2_ of pyridine ring), δ 56.79 ppm (OCH_3_), 124.14(CF3), δ 135.44 (CH imidazole), 113.22-147.46 (Aromatic carbons). MS analysis for C_20_H_16_F_3_N_3_O_3_ Calcd mass 403.11, found (m/z, ESI+) (M^+^ +1): 404.36.

*N-*(3,4-difluorophenyl)-2- [3-(4-fluorobenzoyl)-6,7-dimethoxy -4-oxoquinolin-1-yl] acetamide (**Compound 8)**. ^1^H NMR (DMSO-d6) δ: 3.85 (s, 6H, 2 OCH_3_), 5.23 (s, 2H, CH_2_) 7.02 (s, 1H, <u>Ar</u>H), 7.32 (m, 3H, ArH), 7.45 (q, 1H, ArH), 7.62 (s, 1H, ArH), 7.83 (m, 3H, Ar-H), 8.33 (s, 1H, quinolone 2-CH), 10.80 (s, 1H, NH of acetamide). ^13^C NMR (DMSO-*d6*): δ 56.79 ppm: 2 singlet signals for 2 OCH_3_, δ 35.19 ppm for <u>C</u>H_2_-C=O, δ 104.96-165.72 ppm for aromatic carbons, δ 169.84, 174.96, 188.76 ppm for 3 C=O. MS analysis for C_26_H_19_F_3_N_2_O_5_ Calcd mass 496.12, found (m/z, ESI+) (M^+^ +1): 497.37.

*N*-(4-fluorophenyl) 2- [6,7-dimethoxy-3 -(4-methoxybenzoyl) -4-oxoquinolin-1-yl]-acetamide (**Compound 9)**. ^1^H NMR (DMSO-d6) δ: 3.85 (s, 9H, 3 OCH_3_), 5.24 (s, 2H, CH_2_) 7.02 (m, 3H, <u>Ar</u>H), 7.2 (t, 2H, ArH), 7.60 (m, 3H, ArH), 7.74 (d, 2H, ArH), 8.3 (s, 1H, quinolone 2-CH), 10.62 (s, 1H, NH of acetamide). ^13^C NMR (DMSO-*d6*): δ 53.19-56.79 ppm (3 singlet signals for 3 OCH_3_), δ 56.79 ppm (CH_2_-C=O), δ 104.96-163.49 ppm (aromatic carbons), δ 169.84, 174.56, 188.76 ppm for 3 C=O. MS analysis for C_27_H_23_FN_2_O_6_ Calcd mass 490.48, found (m/z, ESI+) (M^+^ +1): 491.48.

### Computational Methods

Protein and database preparation: The X-ray crystal structure of α/β tubulin in complex with colchicine-DAMA (PDB code: 1SA0) was downloaded from protein data bank https://www.rcsb.org/ and used for virtual screening. Chains C and D were removed, the protein structure was prepared by inserting the missing loop regions using MODELLER^42^ via the UCSF Chimera graphical interface.^43^ Hydrogen atoms were added, and water molecules were removed using MOE. A set comprising the first 100,000 compounds was selected from the clean, druglike subset (9.9 M compounds) of the ZINC15 database to use for virtual screening. The compounds were saved as mdb format files by MOE. A 3D structure-based pharmacophore was constructed from the tubulin-colchicine complex (1SA0) using the Protein-Ligand Interaction Fingerprints (PLIF) application implemented in MOE. The pharmacophore model compromises seven features: three H-bond acceptors and one H-bond donor (F1, F2, F7 and F5) respectively; two hydrophobic centres (F4 and F6), and an aromatic centre (F3). These represent i) the two acceptors F1 and F7 corresponding to interaction with Cysβ241, ii) The third acceptor (F2) corresponding to the interaction of Val α181, iii) aromatic centre (F3), iv) two hydrophobic centres (F4 and F6) corresponding to hydrophobic interaction with Leuβ248, Alaβ250, Leuβ255, Asnβ258, Alaβ316, and Valβl318. The pharmacophore model was employed as a search query using MOE to identify commercial compounds targeting the colchicine binding site, matching at least 4 of the 7 pharmacophore features. This process afforded 2476 compounds for virtual screening by docking described below.

Molecular docking was performed using three programs: BUDE, AutoDock and MOE. Validation of the screening model and applied protocol was carried out by re-docking the native ligand (colchicine) in the binding site of the tubulin protein (1SA0). The RMSD value was then calculated with respect to the co-crystallized ligands. An RMSD value ≤ 1.0 Å between the X-Ray structure and the best-scored conformations of the native ligand, the docking process was considered successful.^44^ The RMSD between the re-docked and co-crystal ligand is less than 1 Å for the three programs, indicating a consensus of the methods and consistency of pose prediction. The predicted binding energies for colchicine re-docked by BUDE, AutoDock and MOE were -100.27 kJ/mol, -9.52 kcal/mol and -5.1 kcal/mol) respectively.

### Virtual screening with BUDE

Molecular docking was performed using the Bristol University Docking Engine (BUDE) for the compounds filtered by the 3D pharmacophore into the tubulin-colchicine binding site. The BUDE search area was defined as grid centred on the native ligand (X=42.848, Y=52.376, Z=-8.531). BUDE is a rapid rigid-body docking program, hence ligand flexibility is achieved by docking multiple conformers of each ligand. All compounds with a predicted binding energy better than colchicine were selected (177 compounds with binding energy ≤ -100 kJ/mol).

### Virtual screening with AutoDock 4.2

The native ligand was removed from the crystal structure (PDB:1SA0) and AutoDock.4.2 used to convert both protein structure and the native ligand separately into PDBQT format. Polar hydrogen atoms and Kollman charges were assigned to the protein. Gasteiger partial charges were assigned to the ligand and non-polar hydrogen atoms merged with their heavy atoms Rotatable bonds in the ligand were defined using an AutoDock utility, AutoTors. The grid box was placed at the centroid of native ligand (X = 42.845, Y = 52.376 and Z = -8.531), the box size was 100 × 100 × 100 Å with a 0.2 A grid spacing and the grid map was calculated using Autogrid tool and saved as a gpf file. Docking was performed using the Lamarckian genetic algorithm, each docking experiment was performed 100 runs, the configuration file was saved as dpf format. Raccoon was used to prepare all the 2476 ligands to perform a docking with Autodock 4.2, the Raccoon software splits the multi-structure files of the ligands to separate PDBQT input files and generate configuration files and scripts for both and Autodock. The results were sorted according to the lowest predicted binding energy. 226 compounds had predicted binding energies ≤ -9.52 kcal/mol (the colchicine binding energy calculated with AutoDock).

### Virtual screening with MOE

The same protein (1SA0) and set of ligands from the 3D pharmacophore filtration and converted to mdb format. MOE was used to add hydrogen atoms to the ligands and energy minimised until the gradient of energy with respect to coordinates fell below 0.05 kcal mol^-1^Å^- 1^ under the was MMFF94X force field. The binding site was defined as the colchicine site. Ligands were docked using the Triangle Matcher method with the London dG scoring function. Refinement was performed using the rescoring affinity dG method. The lowest energy pose was chosen for each docked compound yielding 188 compounds with a binding energy lower than colchicine (−5.1 kcal/mol).

### Selection of compounds for testing

Next the requirement for a compound to be present in at least two docking search results was applied, giving 99 compounds (Supplementary Data File S1.xlsx). These the 99 compounds were re-docked with MOE to allow a consistent set for visualization with Pymol.^39^ Further selection by inspection was performed as described in Results This process gave a shortlist of 13 compounds (Tables S1 and S2). The shortlisted hits were screened for pan assay interference compounds (PAINS) using the online PAINS filters at http://zinc15.docking.org/patterns/home/ and http://www.cbligand.org/PAINS/. The 13 compounds passed both filters. and were purchased from MCULE (USA) for experimental testing.

### Molecular Dynamics (MD) simulations

The recent structure of a tubulin tetramer with a stathmin-4 domain was used as the basis for the MD simulations (6F7C; 2.0 Å) firstly, subunits C and D were removed and stathmin truncated at N91. Crystallographic waters, cofactors GTP and GTP and metal ions associated with subunits A and B were retained apart from waters 653A and 691B at the colchicine site. Ligand coordinates of the six docked complexes of tubulin with colchicine, **6, 8, 9, 14** and **15** were transferred to this model and simulations performed for 3 repeats of 500 ns each using GROMACS 2019.4 and 2021.2 ^46^ as follows. Pdb2gmx was used to add hydrogen atoms to the protein consistent with pH 7 and generate a topology file under the Amber99-SB-ildnb force field.^47^ Acpype^48^ was used to generate topology files of the compounds **6, 8** and **9** under the GAFF force field.^49^ The ligand and protein complexes were centred in a triclinic box with a minimum margin of 1.5 nm and filled with TIP3P water. The system was neutralized by adding sodium and chloride ions to give an ionic strength of 0.15 M. The energy minimization (5000 steps) was conducted using steepest descents. All simulations were performed as NPT ensembles at 310 K under periodic boundary conditions. The Particle Mesh Ewald (PME) method was used for calculating long range electrostatics, and Van der Waals (VdW) interactions. The cut-off distance for the short-range VdW and Coulombic interactions was set to 1.2 nm.^50^ Pressure was controlled by the Parrinello−Rahman barostat and temperature by the V-rescale thermostat. The simulations were integrated with a leap-frog algorithm over a 2 fs time step, constraining h-bond vibrations with the P-LINCS method. Molecular dynamics simulations were carried out for 500 ns on BlueCrystal, the University of Bristol’s high-performance computing machine and the GW4 tier-2 machine Isambard. Simulation analyses were carried out using GROMACS tools, Xmgrace and gnuplot were used for plotting data, molecular graphics manipulations and visualizations were performed using Chimera v1.14^43^, VMD v1.9.4^51^ and OpenPymol v1.8^45^.

## Biological Methods

### Cell culture and maintenance

Breast cancer cell lines including breast cancer cell lines (MCF-7), triple-negative breast cancer (MDA-231), adenocarcinomic human alveolar basal epithelial cells (A549), and normal fibroblast cells (F180) were cultured in Roswell Park Memorial Institute media (RPMI, Sigma-Aldrich, St. Louis, MO, USA) supplemented with 10% fetal bovine serum, and 1% penicillin/streptomycin. All cell lines were purchased from the European Collection of Cell Cultures (ECACC, UK). Cell lines were incubated at 37° C in a humidified incubator containing 5% CO_2_.

### Antiproliferative and cytotoxic activity

The antiproliferative activity of compounds **1-13** were evaluated by testing their effects on the aforementioned cell lines 3-(4, 5-dimethyl thiazolyl-2)-2,5-diphenyltetrazolium bromide (MTT) assay as described before^17^. The cancer cell lines were seeded as 1 × 10^4^ cells per well in 96-well flat bottom plates for 24h. The cells were then treated with 10 μM of compounds **1-13** and incubated for 48h at 37°C in a humidified incubator containing 5% CO_2_. Paclitaxel and 0.1% DMSO were employed as positive and negative controls, respectively. The culture media was then removed, washed, and then incubated with 200 μl culture media containing 0.5 mg/ml MTT for 2 hr. The blue formazan crystals, converted from the yellow MTT by viable cells, were dissolved by adding 200 μl DMSO and measured spectrophotometrically at 570 nm in a microplate reader (Thermo-Scientific, Vantaa, Finland). Cell viability was calculated, using the following formula: % of living cells = (OD experimental) / (OD control) × 100, while % of cell death was calculated by subtracting the living cells from the total number of cells. To determine the IC_50_ of the active compounds, the cells were treated with different concentrations 2.5, 5, 10, 20 μM of compounds (**6, 8** and **9**).

### Immunofluorescence assay

A549 cells (5×10^4^/well) were plated on coverslips in 6-well plates and treated with compounds **6, 8** and **9** at concentrations of 16, 13, and 9 μM, respectively for 24 h. Paclitaxel (14 nM) and Colchicine (0.1 μM) used as positive controls while 0.1% DMSO used as a negative control. The cells were then rinsed twice with PBS, fixed with 3.7% paraformaldehyde, and permeabilized with 0.1% Triton X-100. The cells were blocked with 1% BSA in PBS for 1 h prior to incubation with anti-β-tubulin mouse monoclonal antibody (#86298, Cell Signaling, San Francisco, CA, USA) overnight at 4°C. The cells were washed with PBS for 1 h in the dark, and then incubated with Alexa Fluor® 488 secondary antibodies (Abcam). The cellular microtubules were observed with a fluorescence microscope (Olympus BX43, Japan).

### In vitro tubulin polymerization assay

Tubulin polymerization was analyzed *in vitro* using a Tubulin Polymerization Assay kit (Cytoskeleton, Denver, CO). Briefly, 2 mg/ml Porcine tubulin was dissolved in buffer 1 (80 mM PIPES, 2 mM MgCl_2_, 0.5 mM EGTA pH 6.9, 10 μM fluorescent reporter, 1 mM GTP, 15% glycerol). This solution was transferred to a pre-warmed 96-well plate and treated with 15 μM of the test compounds **6, 8** and **9**. Colchicine and paclitaxel at 3 μM were employed as positive controls, while 0.1% DMSO treated-cells were employed as a negative control. Tubulin polymerization was monitored at 37°C for 60 min using fluorescence microscopy. The reading speed was programmed at 1 cycle/min with excitation and emission wavelengths of 360 and 450 nm, respectively, using the Varioskan Flash spectral scanning multimode reader (Thermo Fisher Scientific). The 100% polymerization value was defined as the area under the curve (AUC) of the untreated control.

### Extraction of soluble and polymerized tubulin fractions

Extraction was done using the protocol previously reported .^52^ After treatment with drugs for 24 h, medium containing cells in suspension was recovered and pooled with adherent cells scraped in PBS pre-warmed at 37°C. After centrifugation 5 min at 400 x g, wash with PBS, cells were extracted for 5 min with pre-warmed at 37°C microtubule-stabilizing buffer (0.1 M PIPES pH 6.9, 14.5% glycerol, 0.5% Triton X-100, 20 mM EGTA and 5 mM MgCl2) containing Complete (Sigma FAST Protease Inhibitor Cocktail Tablet) and 10 ng/ml paclitaxel (Sigma). After centrifugation at 20 000 x g for 10 min at 25°C, supernatants containing soluble fractions were transferred to a new tube, while polymerized fractions in pellets were recovered by incubation in RIPA buffer for 45 min on ice followed by centrifugation for 10 min at 20 000 x g. The total protein content in the samples was determined using DC protein assay kit (Bio-Rad, Hercules, CA, USA). Equivalent aliquots from polymeric fractions were measured using Human Beta-tubulin ELISA Kit (Abcam) following the manufacturer’s instructions.

### Determination of the IC_50_ of compound 6, 8 and 9

The IC_50_ of compounds **6, 8 and 9** required to inhibit tubulin polymerization was measured using ELISA kit (Cat. # BK011P, Cytoskeleton, Denver, CO) with tubulin protein (Cat. #T240-DX, Cytoskeleton, Denver, CO) according to the manufacturer’s instructions.^53^

### Cell Cycle analysis

MCF7 cancer cells were treated with compounds **6** at 6 μM for 24 h. DMSO was employed as vehicle control. The cells were harvested, centrifuged, and the cell pellets were fixed with 70% ethanol on ice for 15 min. The fixed pellets were incubated with propidium iodide (Sigma-Aldrich, St. Louis, MO, USA), staining solution (50 mg/mL PI, 0.1 mg/mL RNaseA, 0.05% Triton X-100) for 1h at room temperature. Cell cycle was assessed by Gallios flow cytometer (Beckman Coulter, Brea, CA, USA).

### Apoptosis assay

The apoptosis assay was carried out using the FITC Annexin-V/PI commercial kit (Becton Dickenson, Franklin Lakes, NJ, USA), MCF-7 cells were treated with compound **6** at 6 μM for 24 hr. 0.1% DMSO was used as negative control. Treated and control cells were stained using V/PI apoptosis kit. Samples were analysed by fluorescence-activated cell sorting (FACS) using flow cytometer run over one hour. Data were analysed using Kaluzav 1.2 (Beckman Coulter).^54^

### Caspase-9 assay

MCF-7 cells were incubated without and with 6 μM of compound **6** for 24 hr. The cells were washed in phosphate buffered saline and cell lysates were collected and level of caspase 9 was determined using ELISA kit (Cat. # EIA-4860, Invitrogen, Carlsbad, CA, USA), and according to manufacturer instruction https://www.thermofisher.com/elisa/product/Caspase-9-Human-ELISA-Kit/BMS2025

### Determination of the effect compound 6 on BAX and Bcl-2 protein levels

MCF-7 cells, which were grown in RPMI1640 containing 10% fetal bovine serum. The cells were treated with compound **6** at 6 μM for 24 h. The cells were then lysed using cell extraction buffer. The collected lysate was diluted in standard diluent buffer and the levels of Bax and Bcl2 were measured as previously reported.^55^

### Statistical analysis

Data were plotted using GraphPad Prism (5.04, La Jolla, CA, USA). A two-way analysis of variance (ANOVA) using Bonferroni’s Multiple Comparison Test was performed and shown as the mean ± SEM of three independent replicates. The statistical significance level was set at *P* < 0.05.

## ASSOCIATED CONTENT

### Supporting Information

ZINC ID lists (Supplementary File S1.xslx). Chemical structure and physical properties of shortlisted compounds and IC_50_ of the promising compounds against cancer cell lines and effect of compound **6** on phases of MCF7 cell cycle (Supplementary Tables S1-S3). Molecular docking and molecular dynamics result (Supplementary Tables S4, S5). Figure S1, schematic view of pharmacophore structure-based virtual screening. Figures S2, S3 clustering of ligands by similarity. Figure S4, RMSD plots of compounds **6, 8, 9, 14** and **15** and colchicine in complex with tubulin during 3 × 500 ns. Figure S5, contact plots of final MD structures. Figures S6-S11 ^1^H and ^13^C NMR and mass spectra of compounds **6, 8, 9**; analytical spectra and data for the short-listed compounds. Simulation structures at 0 ns and 500 ns: protein, ligand and cofactors only (Supplementary File S2.zip).

## Supporting information

revised-ESI

Compound data

Initial and final structures from MD

## Acknowledgments

We thank the Advanced Computing Research Centre at the University of Bristol for provision of High-Performance Computing using BlueCrystal supercomputers. ASFO, DKS and RBS thank BrisSynBio (EPSRC/BBSRC: BB/L01386X/1) for support. We also acknowledge Dr. Esam Rashwan, head of confirmatory diagnostic unit, Vacsera-Egypt, for helping in performing the biological assays. We thank Dr. Amaurys Avila Ibarra for discussions and assistance with software.

## Abstract figure

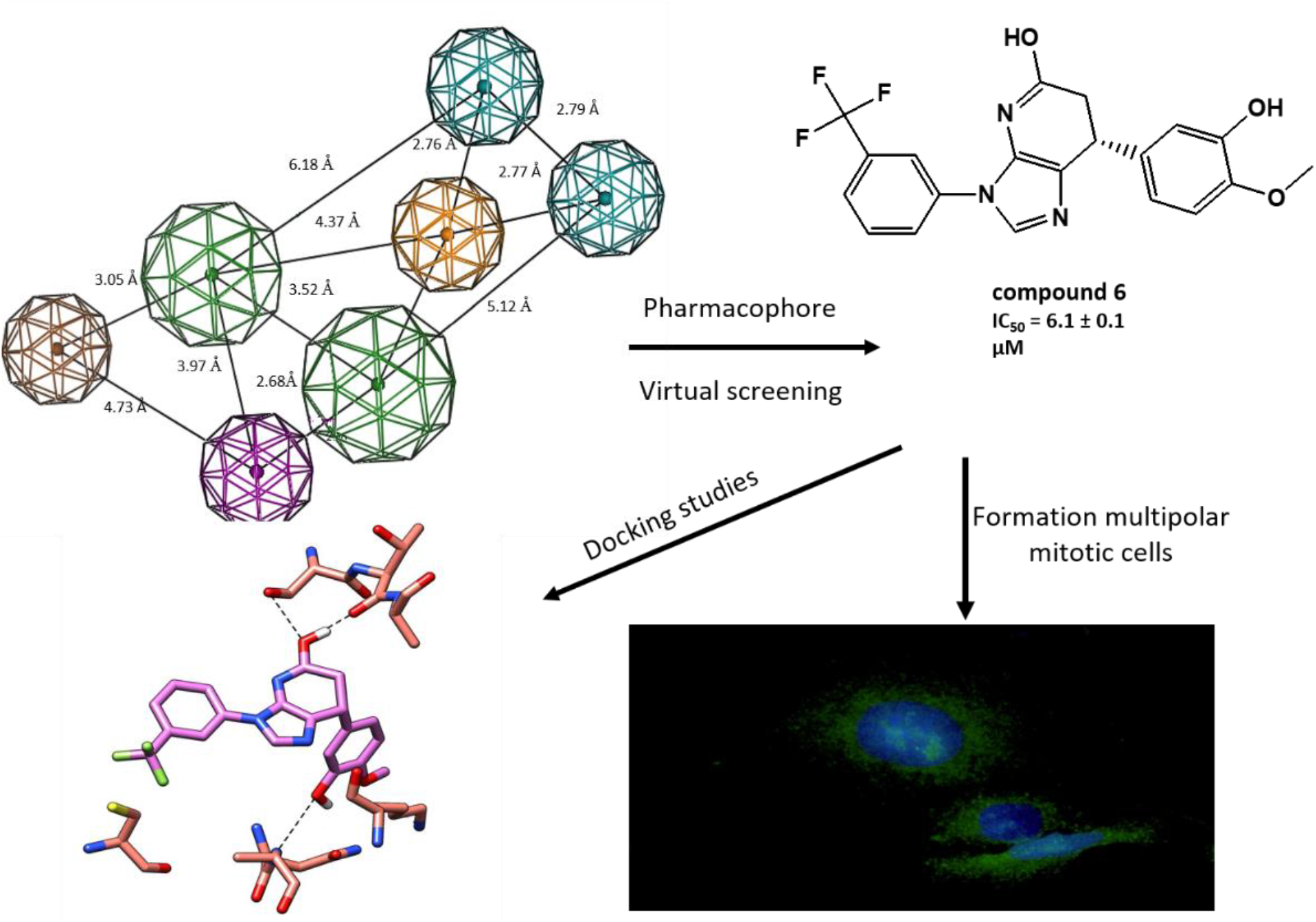

## References

1. Erickson, H. P.; O’Brien, E. T., Microtubule dynamic instability and GTP hydrolysis. Annual review of biophysics and biomolecular structure 1992, 21 (1), 145–166.

2. Prassanawar, S. S.; Panda, D., Tubulin heterogeneity regulates functions and dynamics of microtubules and plays a role in the development of drug resistance in cancer. Biochemical Journal 2019, 476 (9), 1359–1376.

3. Hammond, J. W.; Cai, D.; Verhey, K. J., Tubulin modifications and their cellular functions. Current opinion in cell biology 2008, 20 (1), 71–76.

4. Nogales, E.; Wang, H.-W., Structural intermediates in microtubule assembly and disassembly: how and why? Current opinion in cell biology 2006, 18 (2), 179–184.

5. Mahindroo, N.; Liou, J.-P.; Chang, J.-Y.; Hsieh, H.-P., Antitubulin agents for the treatment of cancer– a medicinal chemistry update. Expert Opinion on Therapeutic Patents 2006, 16 (5), 647–691.

6. Jordan, M. A.; Kamath, K., How do microtubule-targeted drugs work? An overview. Current cancer drug targets 2007, 7 (8), 730–742.

7. Dumontet, C.; Jordan, M. A., Microtubule-binding agents: a dynamic field of cancer therapeutics. Nature reviews Drug discovery 2010, 9 (10), 790–803.

8. Lu, Y.; Chen, J.; Xiao, M.; Li, W.; Miller, D. D., An overview of tubulin inhibitors that interact with the colchicine binding site. Pharmaceutical research 2012, 29 (11), 2943–2971.

9. Tischer, J.; Gergely, F., Anti-mitotic therapies in cancer. Journal of Cell Biology 2019, 218 (1), 10–11

10. Yin, S.; Bhattacharya, R.; Cabral, F., Human mutations that confer paclitaxel resistance. Molecular cancer therapeutics 2010, 9 (2), 327–335.

11. Henriquez, F. L.; Ingram, P. R.; Muench, S. P.; Rice, D. W.; Roberts, C. W., Molecular basis for resistance of Acanthamoeba tubulins to all major classes of antitubulin compounds. Antimicrobial agents and chemotherapy 2008, 52 (3), 1133–1135.

12. McGrogan, B. T.; Gilmartin, B.; Carney, D. N.; McCann, A., Taxanes, microtubules and chemoresistant breast cancer. Biochimica et Biophysica Acta (BBA)-Reviews on Cancer 2008, 1785 (2), 96–132.

13. Li, D. D.; Qin, Y. J.; Zhang, X.; Yin, Y.; Zhu, H. L.; Zhao, L. G., Combined molecular docking, 3D-QSAR, and pharmacophore model: design of novel tubulin polymerization inhibitors by binding to colchicine-binding site. Chemical biology & drug design 2015, 86 (4), 731–745.

14. Lippert III, J. W., Vascular disrupting agents. Bioorganic & medicinal chemistry 2007, 15 (2), 605–615.

15. Field, J. J.; Kanakkanthara, A.; Miller, J. H., Microtubule-targeting agents are clinically successful due to both mitotic and interphase impairment of microtubule function. Bioorganic & medicinal chemistry 2014, 22 (18), 5050–5059.

16. Canela, M.-D.; Pérez-Pérez, M.-J.; Noppen, S.; Sáez-Calvo, G.; Díaz, J. F.; Camarasa, M.-J.; Liekens, S.; Priego, E.-M., Novel colchicine-site binders with a cyclohexanedione scaffold identified through a ligand-based virtual screening approach. Journal of medicinal chemistry 2014, 57 (10), 3924–3938.

17. Lu, Y.; Chen, J.; Wang, J.; Li, C.-M.; Ahn, S.; Barrett, C. M.; Dalton, J. T.; Li, W.; Miller, D. D., Design, synthesis, and biological evaluation of stable colchicine binding site tubulin inhibitors as potential anticancer agents. Journal of medicinal chemistry 2014, 57 (17), 7355–7366.

18. Irwin, J. J.; Sterling, T.; Mysinger, M. M.; Bolstad, E. S.; Coleman, R. G., ZINC: a free tool to discover chemistry for biology. Journal of chemical information and modeling 2012, 52 (7), 1757–1768.

19. Elseginy, S. A.; Lazaro, G.; Nawwar, G. A.; Amin, K. M.; Hiscox, S.; Brancale, A., Computer-aided identification of novel anticancer compounds with a possible dual HER1/HER2 inhibition mechanism. Bioorganic & medicinal chemistry letters 2015, 25 (4), 758–762.

20. Martineau, M.; McIntosh-Smith, S.; Gaudin, W. In Evaluating OpenMP 4.0’s effectiveness as a heterogeneous parallel programming model, 2016 IEEE International Parallel and Distributed Processing Symposium Workshops (IPDPSW), IEEE: 2016; pp 338–347.

21. McIntosh-Smith, S.; Price, J.; Sessions, R. B.; Ibarra, A. A., High performance in silico virtual drug screening on many-core processors. The international journal of high-performance computing applications 2015, 29 (2), 119–134.

22. Morris, G. M.; Huey, R.; Lindstrom, W.; Sanner, M. F.; Belew, R. K.; Goodsell, D. S.; Olson, A. J., AutoDock4 and AutoDockTools4: Automated docking with selective receptor flexibility. Journal of computational chemistry 2009, 30 (16), 2785–2791.

23. Ravelli, R.B.; Gigant, B.; Curmi, P.A., Jourdain, I.; Lachkar, S.; Sorbel, A.; Knossow, M., Insight into Tubulin Regulation from a complex with colchicine and and a stathmin-like domain. Nature 2004, 428, 198–202.

24. Nguyen, T. L.; McGrath, C.; Hermone, A. R.; Burnett, J. C.; Zaharevitz, D. W.; Day, B. W.; Wipf, P.; Hamel, E.; Gussio, R., A common pharmacophore for a diverse set of colchicine site inhibitors using a structure-based approach. Journal of medicinal chemistry 2005, 48 (19), 6107–6116.

25. Massarotti, A.; Coluccia, A.; Silvestri, R.; Sorba, G.; Brancale, A., The tubulin colchicine domain: a molecular modeling perspective. ChemMedChem 2012, 7 (1), 33–42.

26. Dekan, Z.; Sianati, S.; Yousuf, A.; Sutcliffe, K. J.; Gillis, A.; Mallet, C.; Singh, P.; Jin, A. H.; Wang, A. M.; Mohammadi, S. A.; Stewart, M.; Ratnayake, R.; Fontaine, F.; Lacey, E.; Piggott, A. M.; Du, Y. P.; Canals, M.; Sessions, R. B.; Kelly, E.; Capon, R. J.; Alewood, P. F.; Christie, M. J., A tetrapeptide class of biased analgesics from an Australian fungus targets the µ-opioid receptor. Proceedings of the National Academy of Sciences 2019, 116 (44), 22353–22358.

27. Chan, H. H.; Moesser, M. A.; Walters, R. K.; Malla, T. R.; Twidale, R. M.; John, T.; Deeks, H. M.; Johnston-Wood, T.; Mikhailov, V.; Sessions, R. B.; Dawson, W.; Salah, E.; Lukacik, P.; Strain-Damerell, C.; Owen, C. D.; Nakajima, T.; Swiderek, K.; Lodola, A.; Moliner, V.; Glowacki, D. R.; Walsh, M. A.; Schofield, C. J.; Genovese, L.; Shoemark, D. K.; Mulholland, A. J.; Duarte, F.; Morris, G. M., Discovery of SARS-CoV-2 Mpro Peptide Inhibitors from Modelling Substrate and Ligand Binding. bioRxiv 2021

28. Smith, S. A.; Sessions, R. B.; Shoemark, D. K.; Williams, C.; Ebrahimighaei, R.; McNeill, M. C.; Crump, M. P.; McKay, T. R.; Harris, G.; Newby, A. C., Bond, M., Antiproliferative and antimigratory effects of a novel YAP–TEAD interaction inhibitor identified using in silico molecular docking. Journal of medicinal chemistry 2019, 62 (3), 1291–1305.

29. Anukanon, S.; Pongpamorn, P.; Tiyabhorn, W.; Chatwichien, J.; Niwetmarin, W.; Sessions, R. B.; Ruchirawat, S.; Thasana, N., In Silico-Guided Rational Drug Design and Semi-synthesis of C (2)-Functionalized Huperzine-A Derivatives as Acetylcholinesterase Inhibitors. ACS omega 2021, 6 (30), 19924–19939.

30. Cho, S-D.; Kweon, D-H.; Kang, Y-J.; Lee, S-G; Lee, W.S.; Yoon, Y-J., Synthesis of 6,7-dimethoxy-1-halobenzyl-1,2,3,4-tetrahydroisoquinolines. J. Heterocyclic Chem. 1999, 36 (5), 1151–1156.

31. Wang, J.; Miller, D.D.; Li, W., Molecular interactions at the colchicine binding site in tubulin: An X-ray crystallography perspective. Drug Disc. Today 2022 27 (3) 759–776.

32. Korff, M. von, Freyss, J., Sander, T., Flexophore, a New Versatile 3D Pharmacophore Descriptor That Considers Molecular Flexibility. J. Chem. Inf. Model. 2008, 48, (4), 797–810.

33. Supino, R., In vitro toxicity testing protocols. Methods in molecular biology 1995, 43 (2), 137–149.

34. Mooberry, S. L.; Weiderhold, K. N.; Dakshanamurthy, S.; Hamel, E.; Banner, E. J.; Kharlamova, A.; Hempel, J.; Gupton, J. T.; Brown, M. L., Identification and characterization of a new tubulin-binding tetrasubstituted brominated pyrrole. Molecular pharmacology 2007, 72 (1), 132–140.

35. Shelanski, M. L.; Gaskin, F.; Cantor, C. R., Microtubule assembly in the absence of added nucleotides. Proceedings of the National Academy of Sciences 1973, 70 (3), 765–768.

36. Lee, J. C.; Timasheff, S. N., In vitro reconstitution of calf brain microtubules: effects of solution variables. Biochemistry 1977, 16 (8), 1754–1764.

37. Aubry, J. P.; Blaecke, A.; Lecoanet-Henchoz, S.; Jeannin, P.; Herbault, N.; Caron, G.; Moine, V.; Bonnefoy, J. Y., Annexin V used for measuring apoptosis in the early events of cellular cytotoxicity. Cytometry: The Journal of the International Society for Analytical Cytology 1999, 37 (3), 197–204.

38. Parrish, A. B.; Freel, C. D.; Kornbluth, S., Cellular mechanisms controlling caspase activation and function. Cold Spring Harbor perspectives in biology 2013, 5 (6), a008672.

39. Syam, Y. M.; Anwar, M. M.; Kotb, E. R.; Elseginy, S. A.; Awad, H. M.; Awad, G. E., Development of Promising Thiopyrimidine-Based Anti-cancer and Antimicrobial Agents: Synthesis and QSAR Analysis. Mini Reviews in Medicinal Chemistry 2019, 19 (15), 1255–1275.

40. Forbes, C. R.; Sinha, S. K.; Ganguly, H. K.; Bai, S.; Yap, G. P.; Patel, S.; Zondlo, N. J., Insights into thiol–aromatic interactions: A stereoelectronic basis for S–H/π interactions. Journal of the American Chemical Society 2017, 139 (5), 1842–1855.

41. Lichitsky, B.V.; Komogortsev, A.N.; Dudinov, A.A.; Krayushkin, M.M., Three-component condensation of 5-aminoimidazole derivatives with aldehydes and Meldrum’s acid. Synthesis of 3,4,6,7-tetrahydroimidazo[4,5-b]pyridine-5-ones. Russian Chemical Bulletin, Int. Ed., 2012, 61 (8), 1591–1595.

42. Eswar, N.; Webb, B.; Marti-Renom, M. A.; Madhusudhan, M.; Eramian, D.; Shen, M. y.; Pieper, U.; Sali, A., Comparative protein structure modeling using Modeller. Current protocols in bioinformatics 2006, 15 (1), 5.6. 1-5.6. 30.

43. Pettersen, E. F.; Goddard, T. D.; Huang, C. C.; Couch, G. S.; Greenblatt, D. M.; Meng, E. C.; Ferrin, T. E., UCSF Chimera—a visualization system for exploratory research and analysis. Journal of computational chemistry 2004, 25 (13), 1605–1612.

44. Pagadala, N. S.; Syed, K.; Tuszynski, J., Software for molecular docking: a review. Biophysical reviews 2017, 9 (2), 91–102.

45. DeLano, W. L., The PyMOL user’s manual. DeLano Scientific, San Carlos, CA 2002, 452.

46. Berendsen, H. J.; Postma, J. v.; van Gunsteren, W. F.; DiNola, A.; Haak, J. R., Molecular dynamics with coupling to an external bath. The Journal of chemical physics 1984, 81 (8), 3684–3690.

47. Lindorff-Larsen, K.; Piana, S.; Palmo, K.; Maragakis, P.; Klepeis, J. L.; Dror, R. O.; Shaw, D. E., Improved side-chain torsion potentials for the Amber ff99SB protein force field. Proteins: Structure, Function, and Bioinformatics 2010, 78 (8), 1950–1958.

48. Da Silva, A. W. S.; Vranken, W. F., ACPYPE-Antechamber python parser interface. BMC research notes 2012, 5 (1), 1–8.

49. Wang, W. R.; Wolf, R.J.; Caldwell, JW; Kollman, PA; Case. DA Journal of Computational Chemistry 2004, 25, 1157–1174.

50. Wang, H.; Dommert, F.; Holm, C., Optimizing working parameters of the smooth particle mesh Ewald algorithm in terms of accuracy and efficiency. The Journal of chemical physics 2010, 133 (3), 034117.

51. Humphrey, W.; Dalke, A.; Schulten, K., VMD: visual molecular dynamics. Journal of molecular graphics 1996, 14 (1), 33–38.

52. Martel-Frachet, V.; Keramidas, M.; Nurisso, A.; DeBonis, S.; Rome, C.; Coll, J.-L.; Boumendjel, A.; Skoufias, D. A.; Ronot, X., IPP51, a chalcone acting as a microtubule inhibitor with in vivo antitumor activity against bladder carcinoma. Oncotarget 2015, 6 (16), 14669.

53. Liliom, K.; Lehotzky, A.; Molnar, A.; Ovadi, J., Characterization of tubulin-alkaloid interactions by enzyme-linked immunosorbent assay. Analytical biochemistry 1995, 228 (1), 18–26.

54. Shahali, A.; Ghanadian, M.; Jafari, S. M.; Aghaei, M., Mitochondrial and caspase pathways are involved in the induction of apoptosis by nardosinen in MCF-7 breast cancer cell line. Research in pharmaceutical sciences 2018, 13 (1), 12.

55. Brambilla, E.; Negoescu, A.; Gazzeri, S.; Lantuejoul, S.; Moro, D.; Brambilla, C.; Coll, J.-L., Apoptosis-related factors p53, Bcl2, and Bax in neuroendocrine lung tumors. The American journal of pathology 1996, 149 (6), 1941.

## References

24. Dlugosz, P. J.; Billen, L. P.; Annis, M. G.; Zhu, W.; Zhang, Z.; Lin, J.; Leber, B.; Andrews, D. W., Bcl-2 changes conformation to inhibit Bax oligomerization. The EMBO journal 2006, 25 (11), 2287–2296.

