## Supplementary material for "Identification and Validation of Novel Microtubule Suppressors with an Imidazopyridine Scaffold through Structure-Based Virtual Screening and Docking": revised-ESI

**ELECTRONIC SUPPORTING INFORMATION**

**Table of Contents**

Table S1: Chemical structures and docking energies of shortlisted compounds.

Table S2: Physical properties of shortlisted compounds.

Table S3: effect of compound 6 on phases of the MCF7 cell cycle.

Table S4: Molecular docking results for compounds **6**, **8** and **9**.

Table S5: RMSD averages of the molecular dynamics simulations.

Figure S1: Schematic view of pharmacophore structure-based virtual screening.

Figure S2: Clustering of the best 61 compounds.

Figure S3: Clustering of the best 61 compounds and 50 ligands from complexes of known structure.

Figure S4: RMSD plots of the molecular dynamics simulations.

Figure S5: Ligplot^+^ protein-ligand contact diagrams.

Figures S6-S11: Compound characterisation data, ^1^H, ^13^C NMR, GCMS.

**Table S1:** Chemical structure of shortlisted compounds and their binding energies by BUDE, AutoDock, and MOE.

| Cp. | ZINC-ID | Structure | BUDE binding energy  kJ/mol | AutoDock binding energy  Kcal/mol | MOE  Binding  Energy  kcal/mol |
| --- | --- | --- | --- | --- | --- |
| 1 | Zinc02843810 | 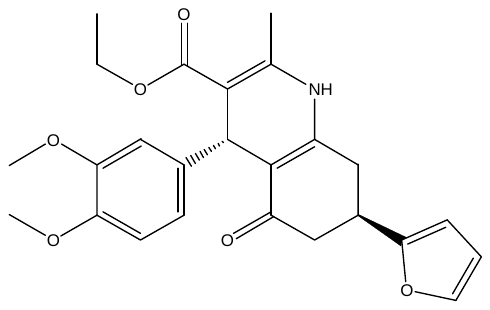 | -111.013 | -9.79 | -6.28 |
| 2 | Zinc02691641 | 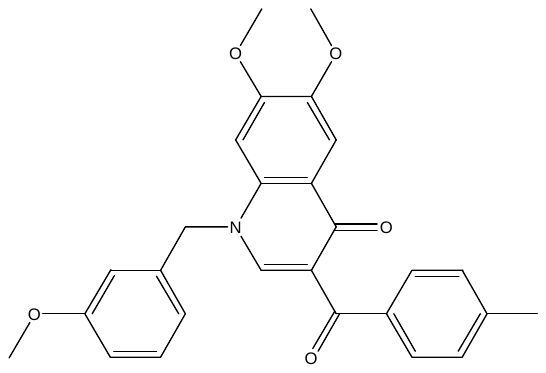 | -92.82 | -10.31 | -7.2 |
| 3 | Zinc03614688 | 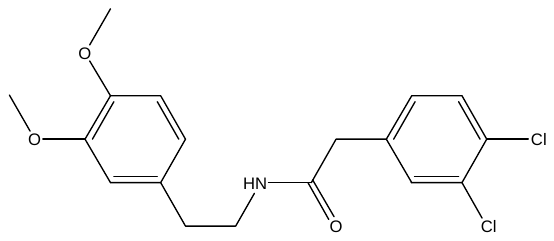 | -91.78 | -7.87 | -5.2 |
| 4 | Zinc49543397 | 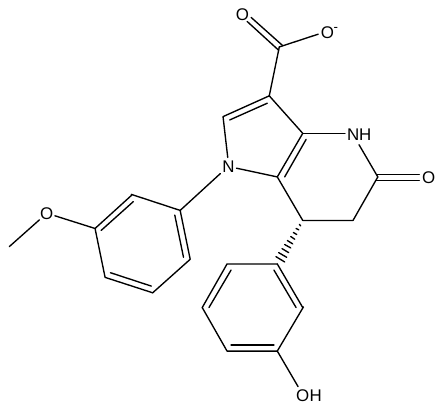 | -104.3 | -9.14 | -8.01 |
| 5 | Zinc02690781 | 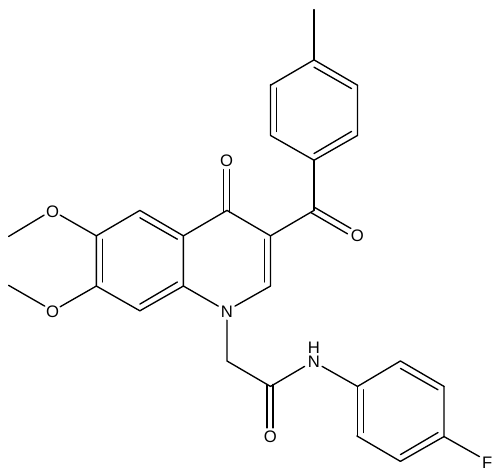 | -100.75 | -10.08 | -8.24 |
| 6 | Zinc36360243 | 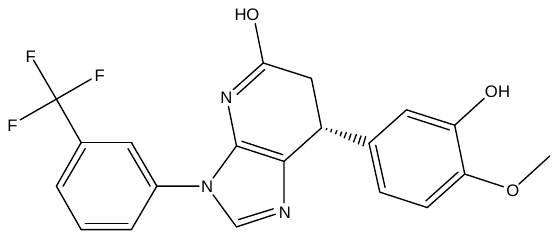 | -100.9 | -8.57 | -10.88 |
| 7 | Zinc18200970 | 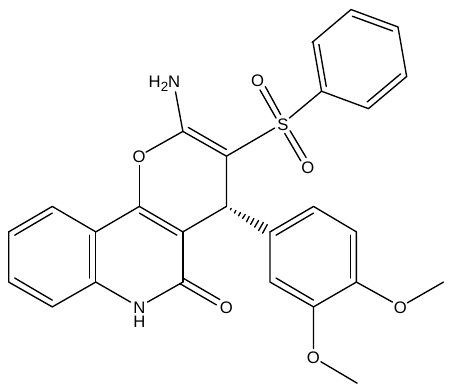 | -125.35 | -9.7 | -6.9 |
| 8 | Zinc02690805 | 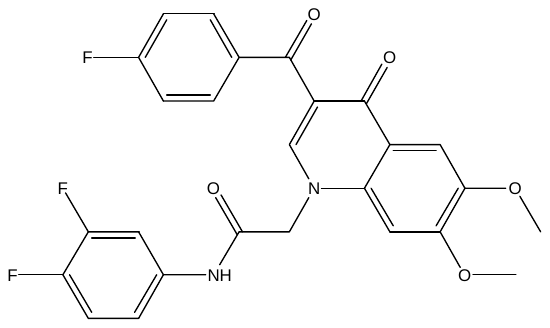 | -119.48 | -9.65 | -9.61 |
| 9 | Zinc02690789 | 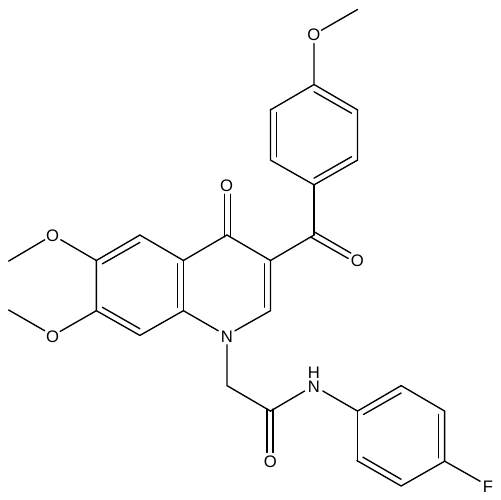 | -108.21 | -10.02 | -8.93 |
| 10 | Zinc07095120 | 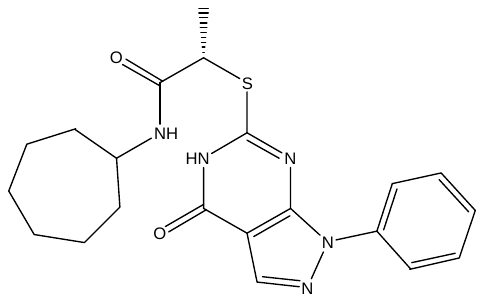 | -83.65 | -9.79 | -5.5 |
| 11 | Zinc11112053 | 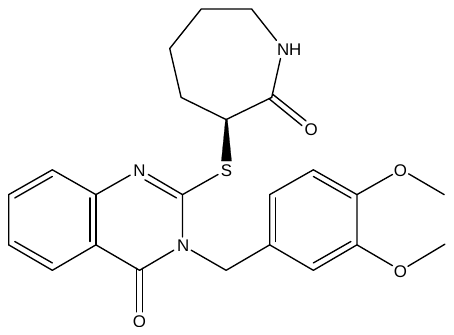 | -83.66 | -9.9 | -6.32 |
| 12 | Zinc23483881 | 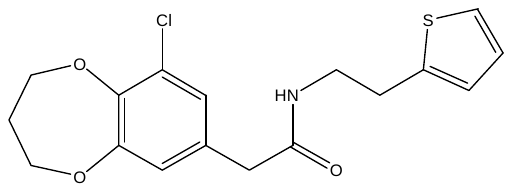 | -84.55 | -9.9 | -5.7 |
| 13 | Zinc03254098 | 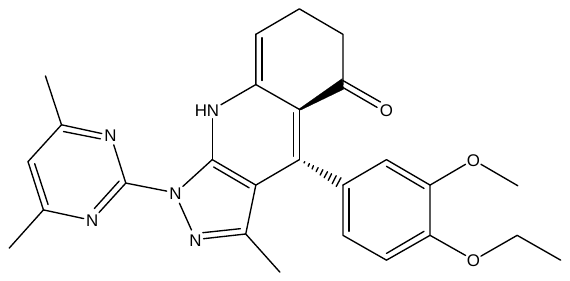 | --105.95 | -10.91 | -6.36 |

**Table S2:** Physical properties of shortlisted compounds calculated by MOE.

| **Compounds** | **ZINC ID** | **MwSt** | **Log p**  **(O/W)** | **H-bond**  **donors** | **H-bond acceptors** | **tPSA** | **Rotatable bond** |
| --- | --- | --- | --- | --- | --- | --- | --- |
| 1 | Zinc02843810 | 437.49 | 2.6 | 1 | 4 | 87.00 | 7 |
| 2 | Zinc02691641 | 443.49 | 4.6 |  | 5 | 65.07 | 5 |
| 3 | Zinc03614688 | 368.26 | 4.08 | 1 | 3 | 47.56 | 3 |
| 4 | Zinc49543397 | 377.37 | 3.16 | 2 | 3 | 103.62 | 4 |
| 5 | Zinc02690781 | 474.48 | 3.79 | 1 | 5 | 84.94 | 8 |
| 6 | Zinc36360243 | 403.36 | 3.9 | 2 | 5 | 79.87 | 4 |
| 7 | Zinc18200970 | 490.53 | 3.4 | 2 | 5 | 116.95 | 5 |
| 8 | Zinc02690805 | 496.44 | 3.8 | 1 | 5 | 84.94 | 8 |
| 9 | Zinc02690789 | 490.48 | 3.4 | 1 | 6 | 94.17 | 9 |
| 10 | Zinc07095120 | 411.53 | 4.05 | 2 | 4 | 88.38 | 6 |
| 11 | Zinc11112053 | 439.53 | 3.7 | 1 | 5 | 80.23 | 6 |
| 12 | Zinc23483881 | 351.85 | 3.3 | 1 | 3 | 47.56 | 6 |
| 13 | Zinc03254098 | 457.53 | 3.2 | 1 | 6 | 91.16 | 5 |

**Table S3** The effect of compound **6** on the phases of the MCF7 cell cycle.

| compounds | %G0-G1 | %S | %G2-M | %Pre G1 |
| --- | --- | --- | --- | --- |
| Control | 53.91 | 42.71 | 3.38 | 1.48 |
| **6** | 36.28 | 29.66 | 34.06 | 27.36 |

**Table S4** Hydrogen bonding of compounds **6,8** and **9** docked into the colchicine binding site.

| Cp | Interacting moiety  in compound | Amino acid involved | Distance  Å | Type of interaction |
| --- | --- | --- | --- | --- |
| **6** | OH-imidazopyridine  OH-phenyl | C=O Thrα179  NH Alaβ250 | 2.5  3.1 | H-bond  H-bond |
| **8** | C=O  C=O | SH Cysβ241  OH Serα178 | 3.1  2.9 | H-bond  H-bond |
| **9** | OCH_3_  C=O | SH Cysβ241  OH Serα178 | 3.6  3.0 | H-bond  H-bond |

**Table S5** Average RMSD values (nm) of complexes of **6, 8, 9, 14, 15** and colchicine with tubulin over 1.5 µs of simulation.

| nm | **6** | **8** | **9** | **14** | **15** | **colchicine** |
| --- | --- | --- | --- | --- | --- | --- |
| Average RMSD ligand | 0.14 ± 0.03 | 0.34 ± 0.05 | 0.22 ± 0.08 | 0.22 ± 0.08 | 0.17 ± 0.03 | 0.15 ± 0.04 |
| Average RMSD protein (AB) | 0.25 ± 0.02 | 0.25 ± 0.02 | 0.24 ± 0.02 | 0.26 ± 0.03 | 0.26 ± 0.03 | 0.25 ± 0.02 |

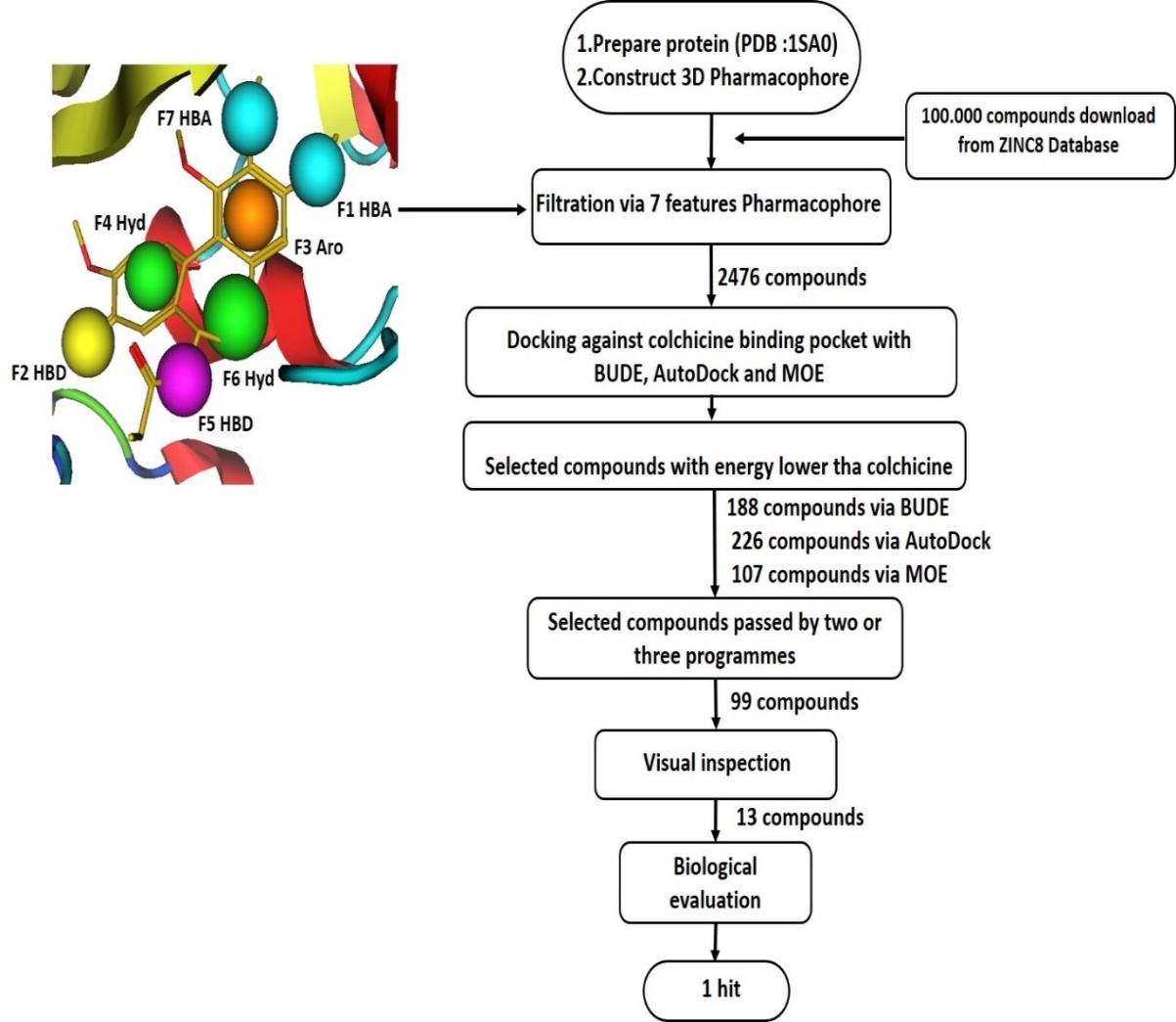

**Figure S1:** Schematic view of pharmacophore structure-based virtual screening.

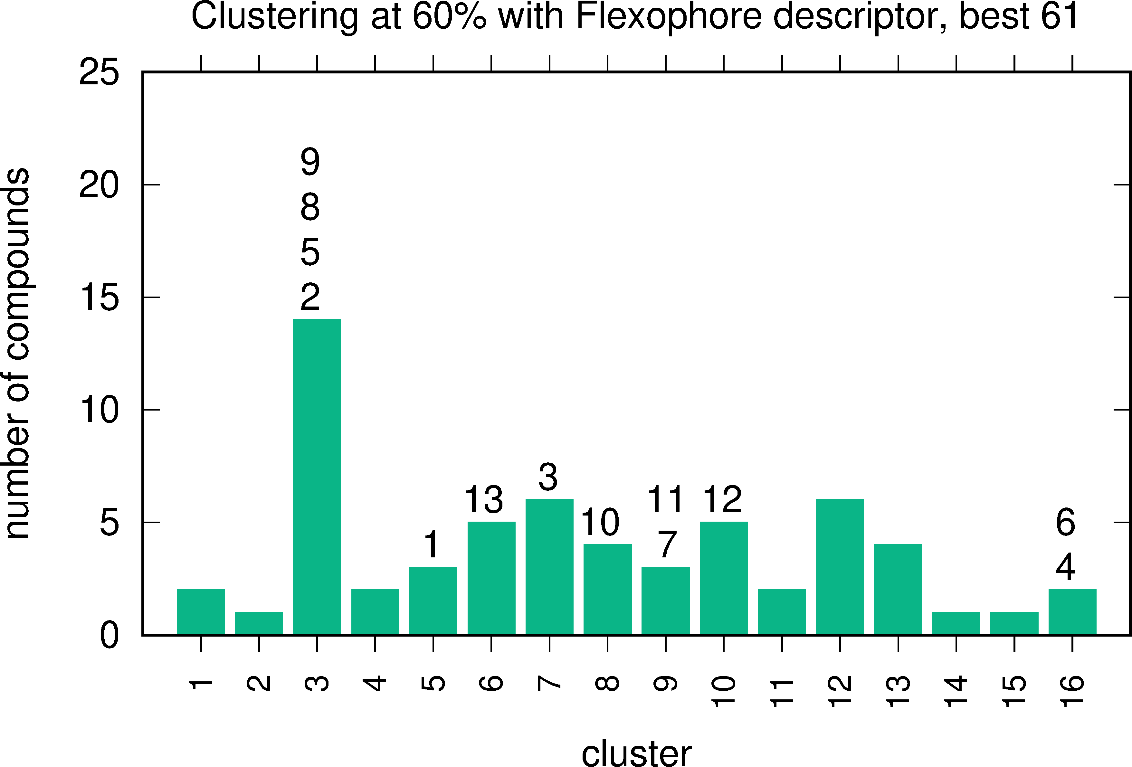

**Figure S2** Clustering of the best 61 compounds with the Flexophore descriptor at 60% similarity. The cluster locations of **1-13** are indicated.

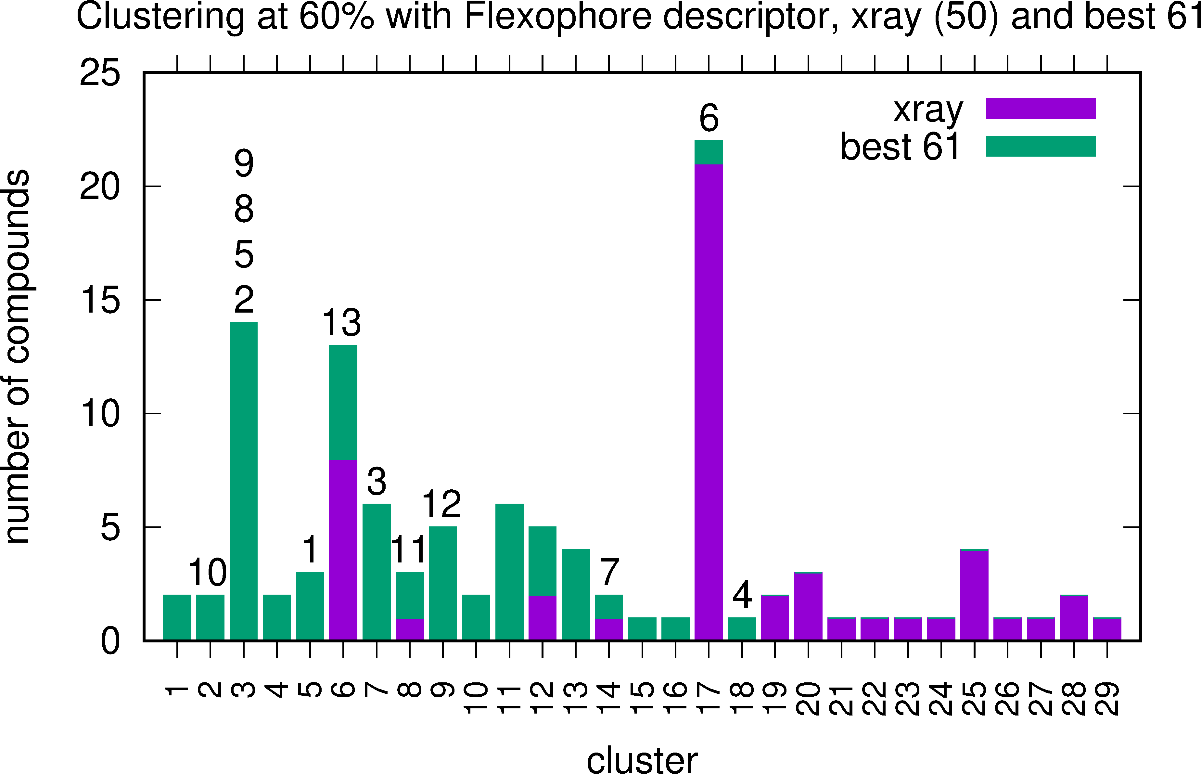

**Figure S3** Clustering of the best 61 compounds with the 50 ligands in the colchicine site from crystal structures reported to date, using the Flexophore descriptor at 60% similarity. The cluster locations of **1-13** are indicated.

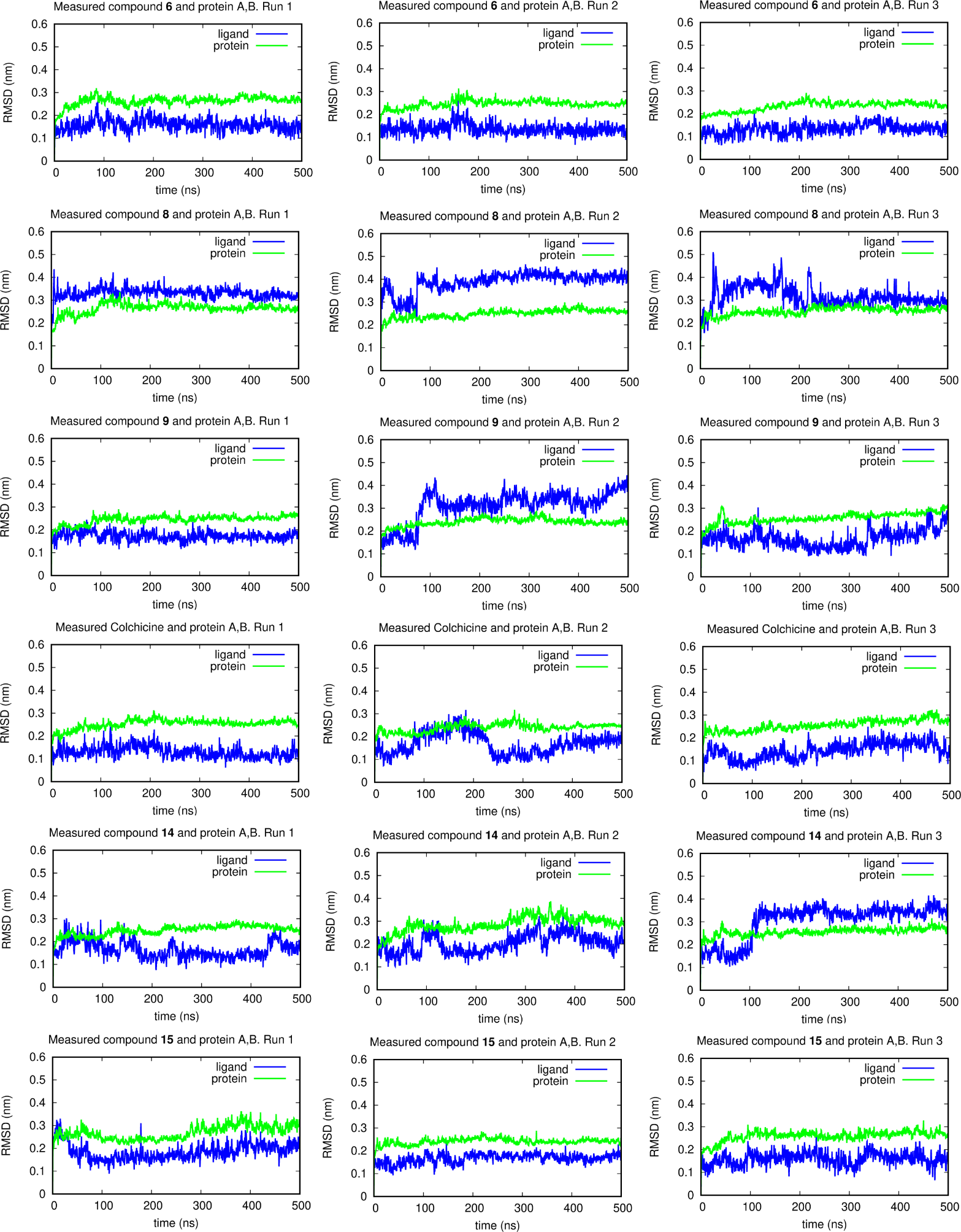

**Figure S4** RMSD plots of all protein atoms of tubulin subunits A and B superimposed on the time = 0 ns structure over the trajectories and measuring all atoms of protein A and B (green) and all atoms of ligand (blue).

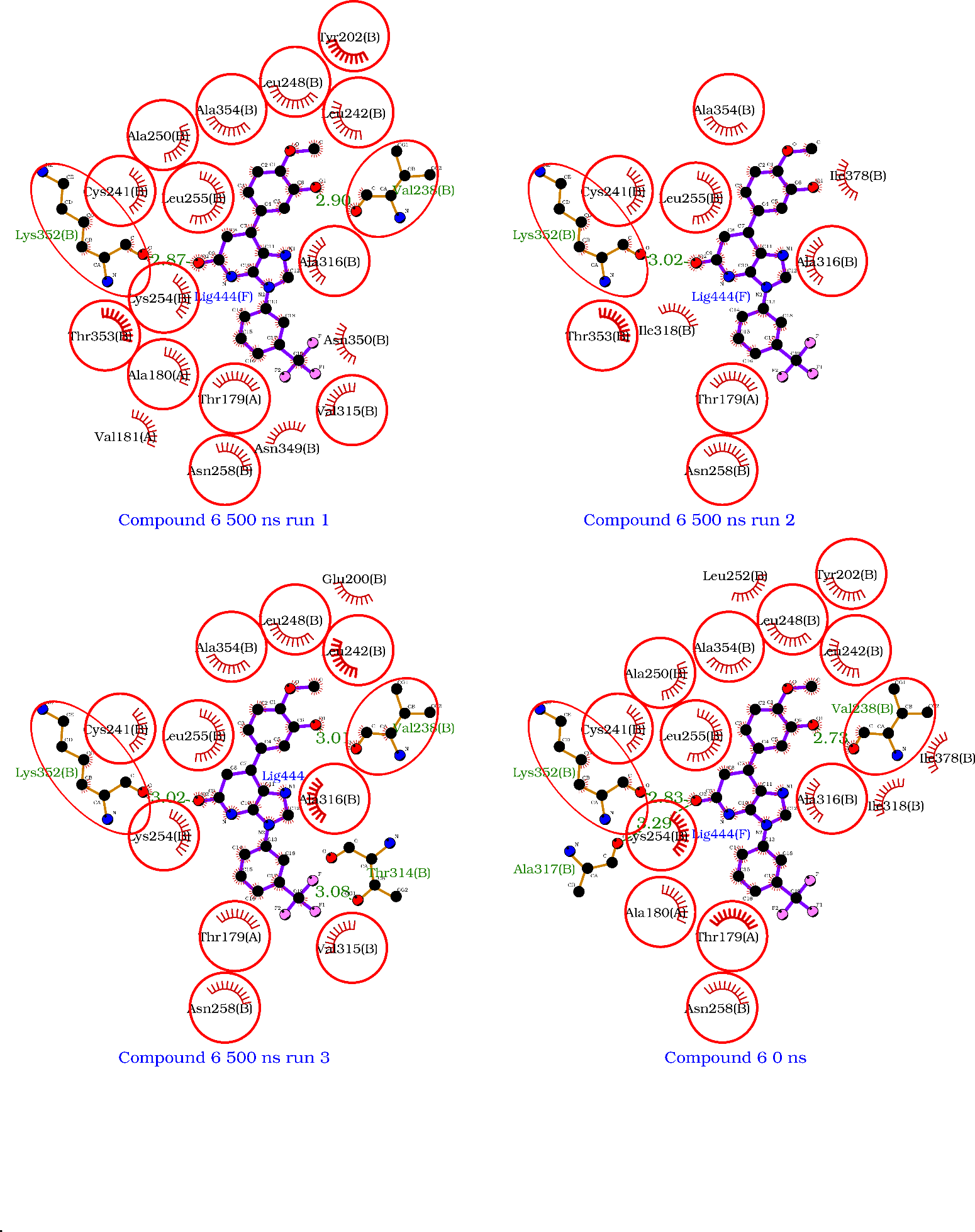

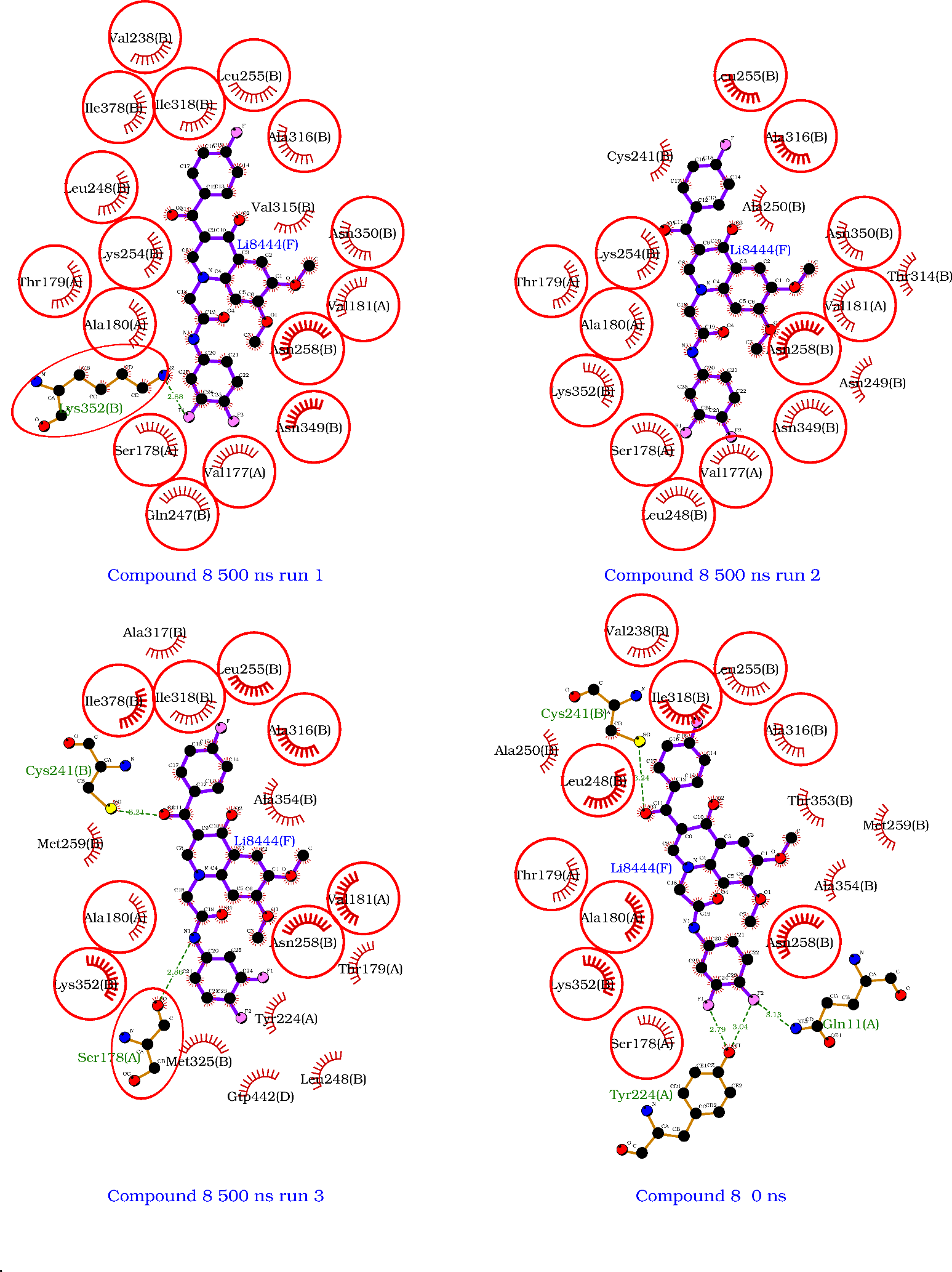

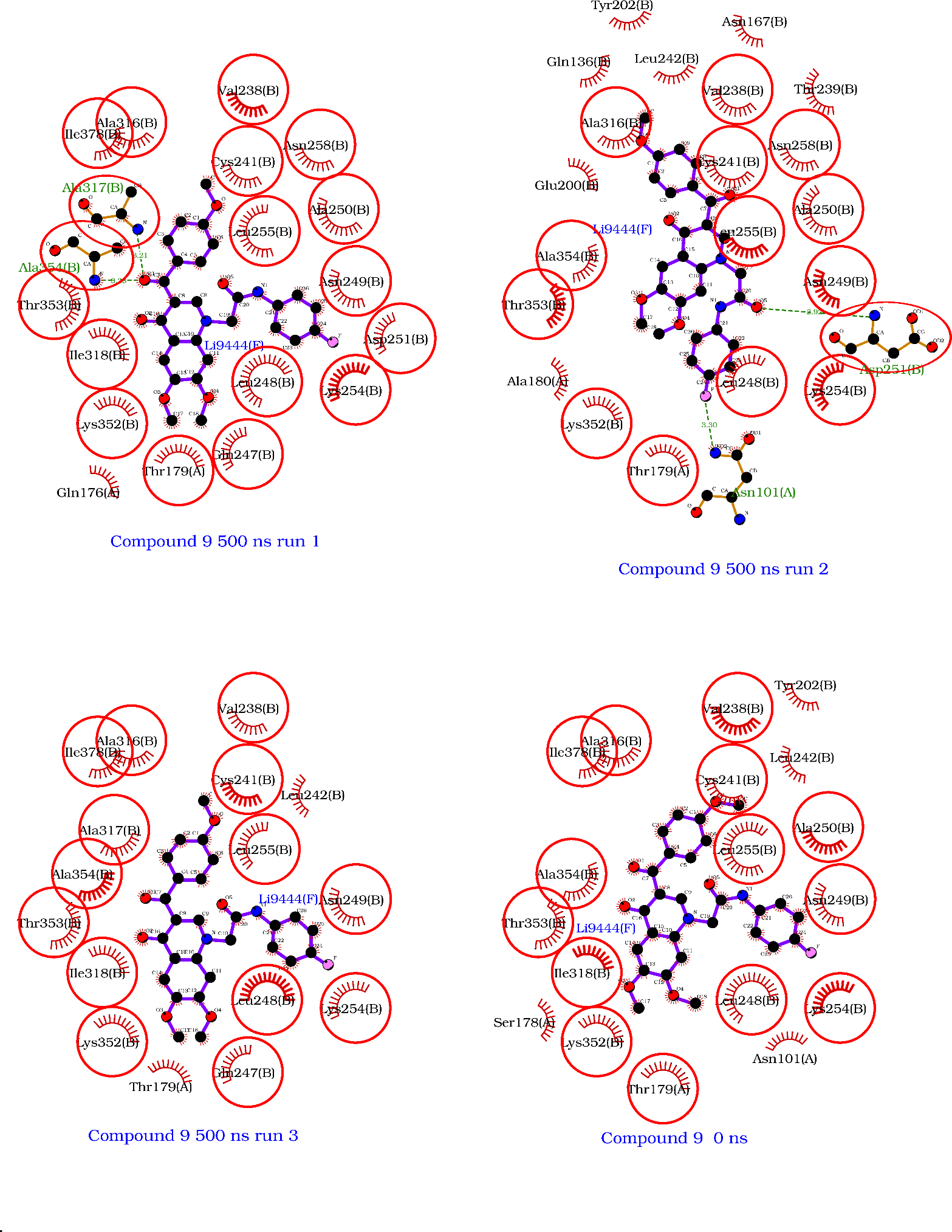

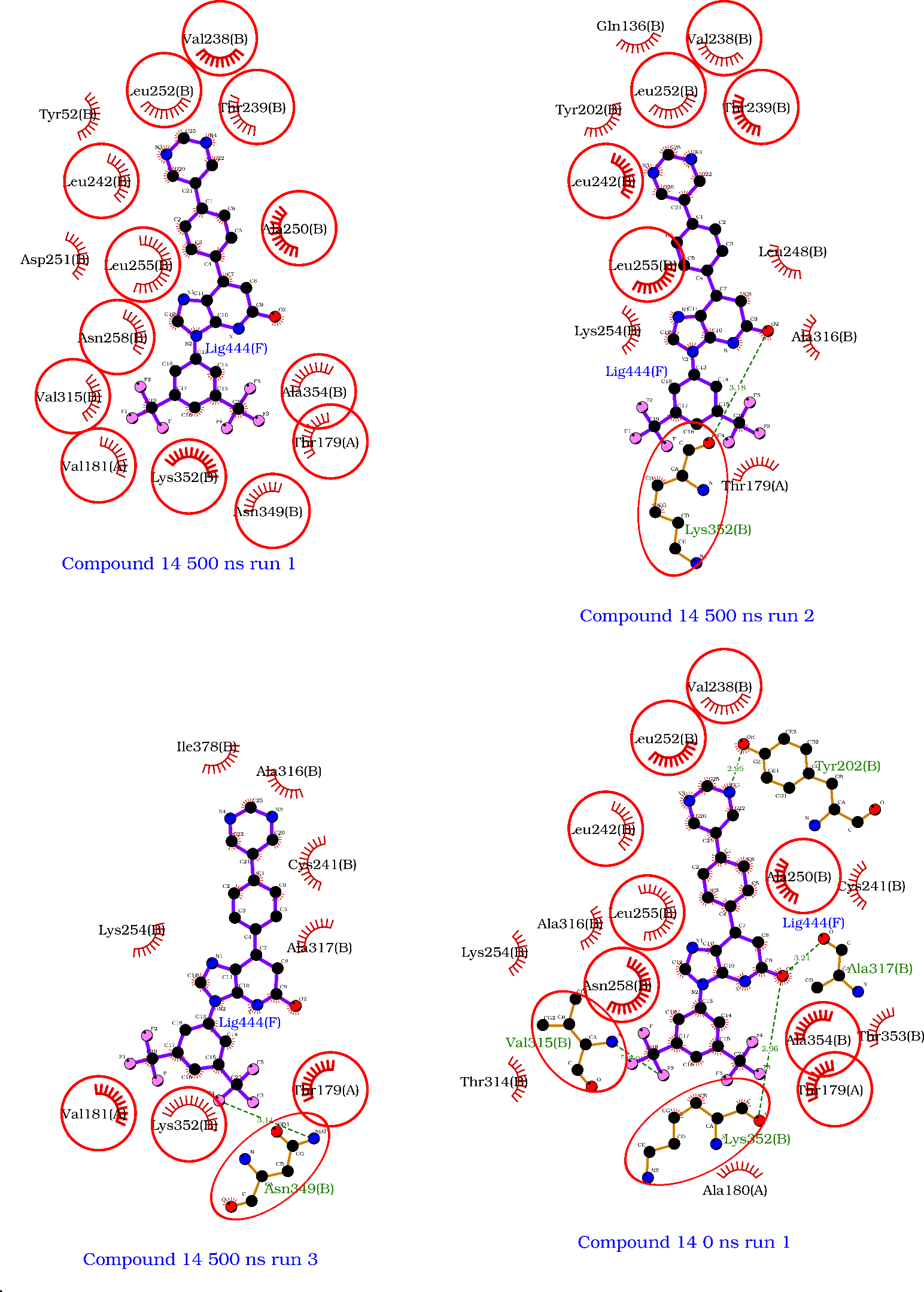

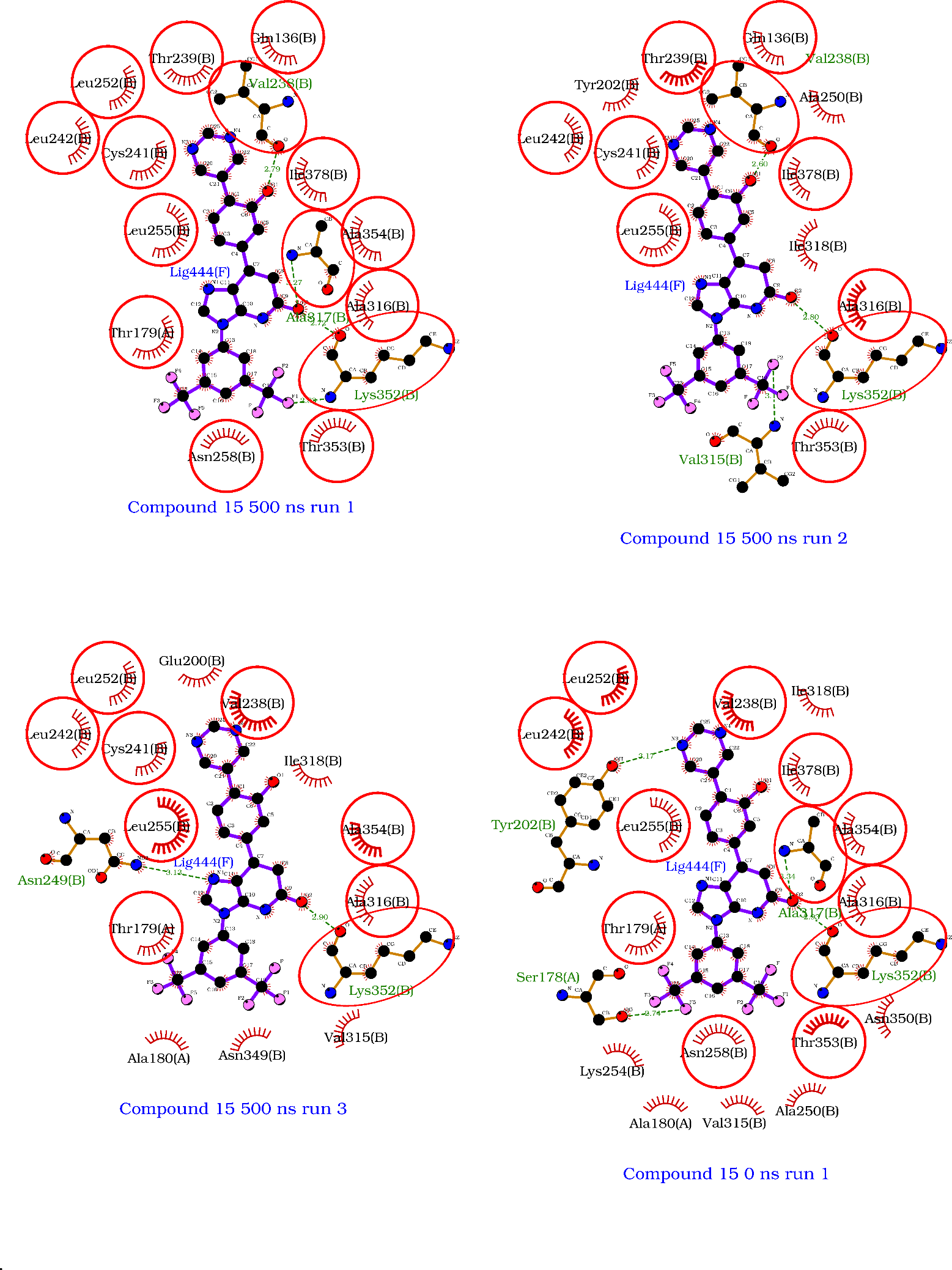

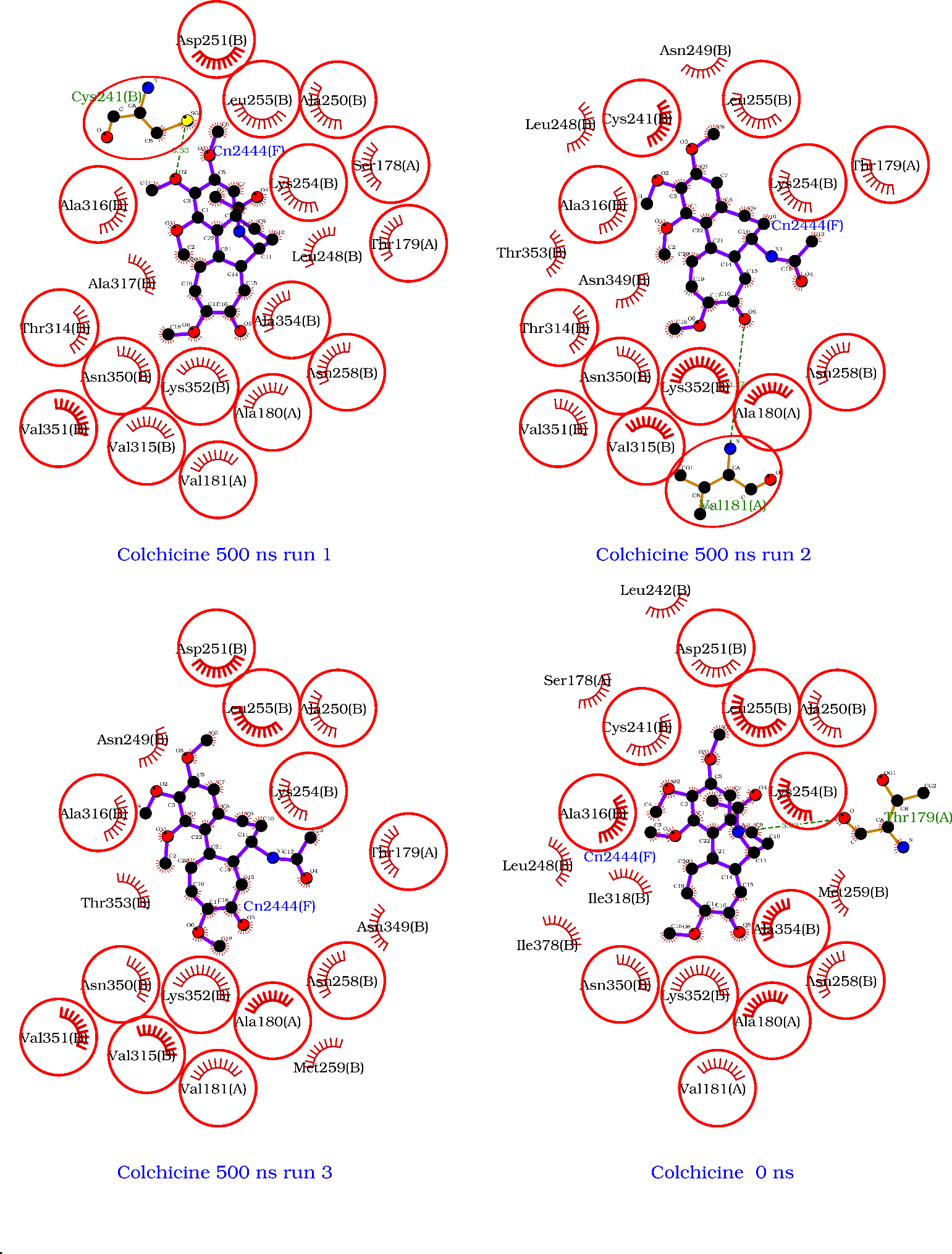

**Figure S5** Ligplot^+^ diagrams of the 500 ns structures for each simulation showing ligand-residue contacts at that moment.

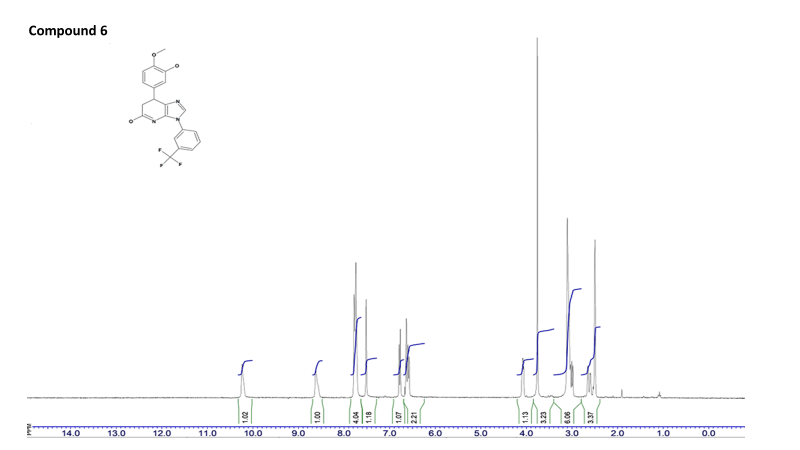

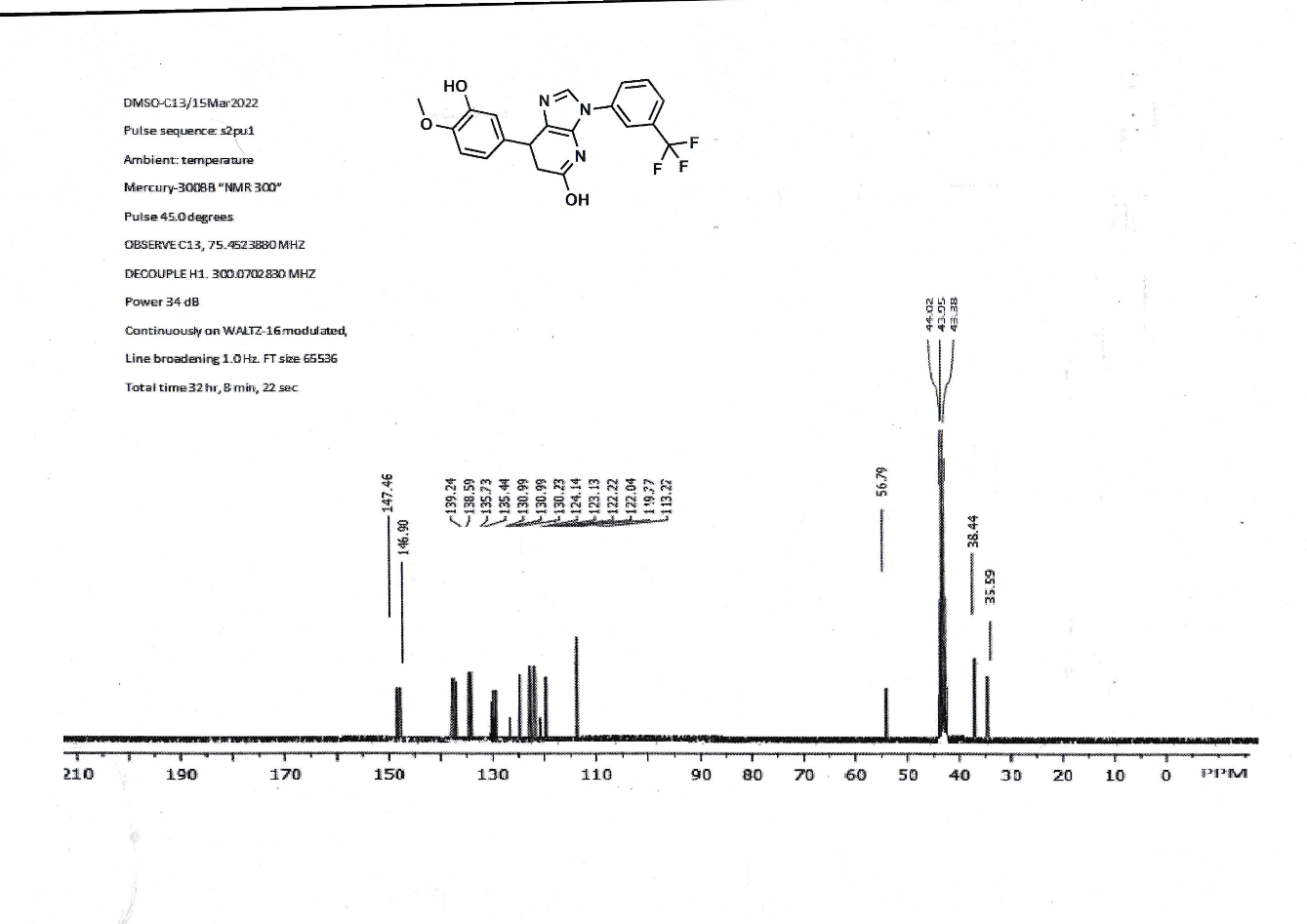

**Figure S6** ^1^H and ^13^C NMR spectra of compound **6**.

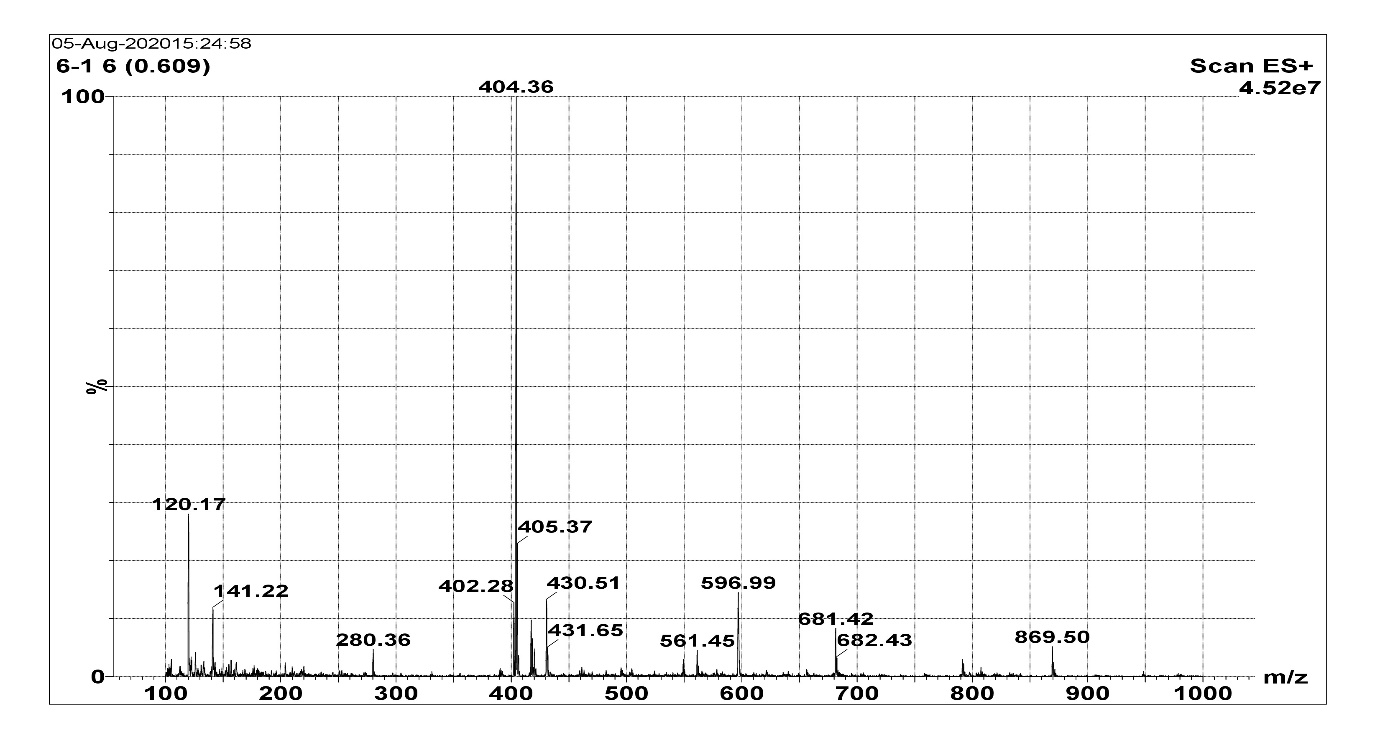

**Figure S7** Mass spectrum of compound 6.

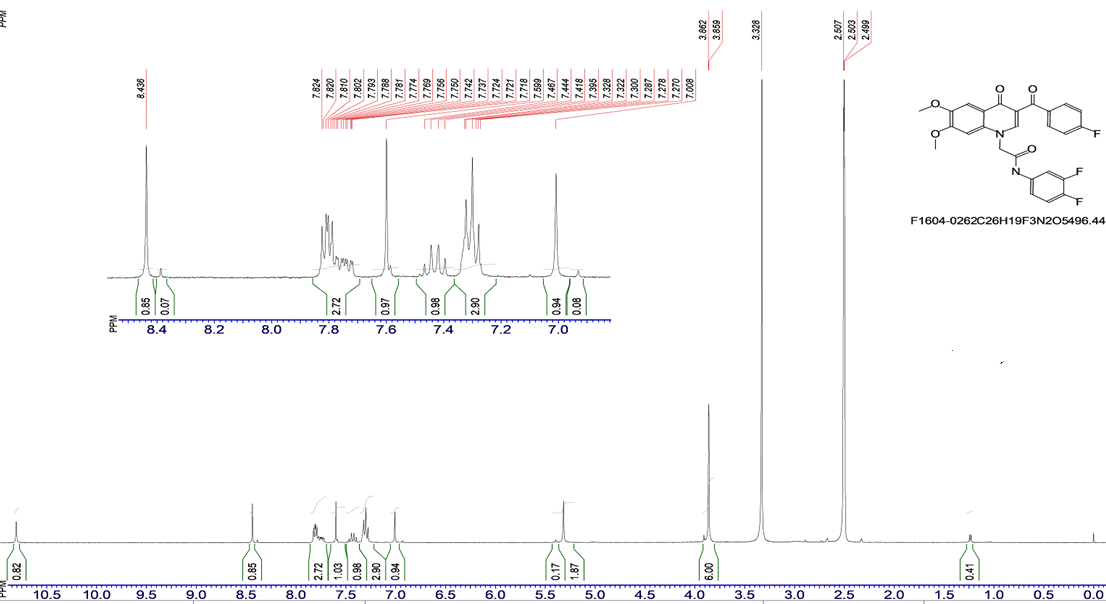

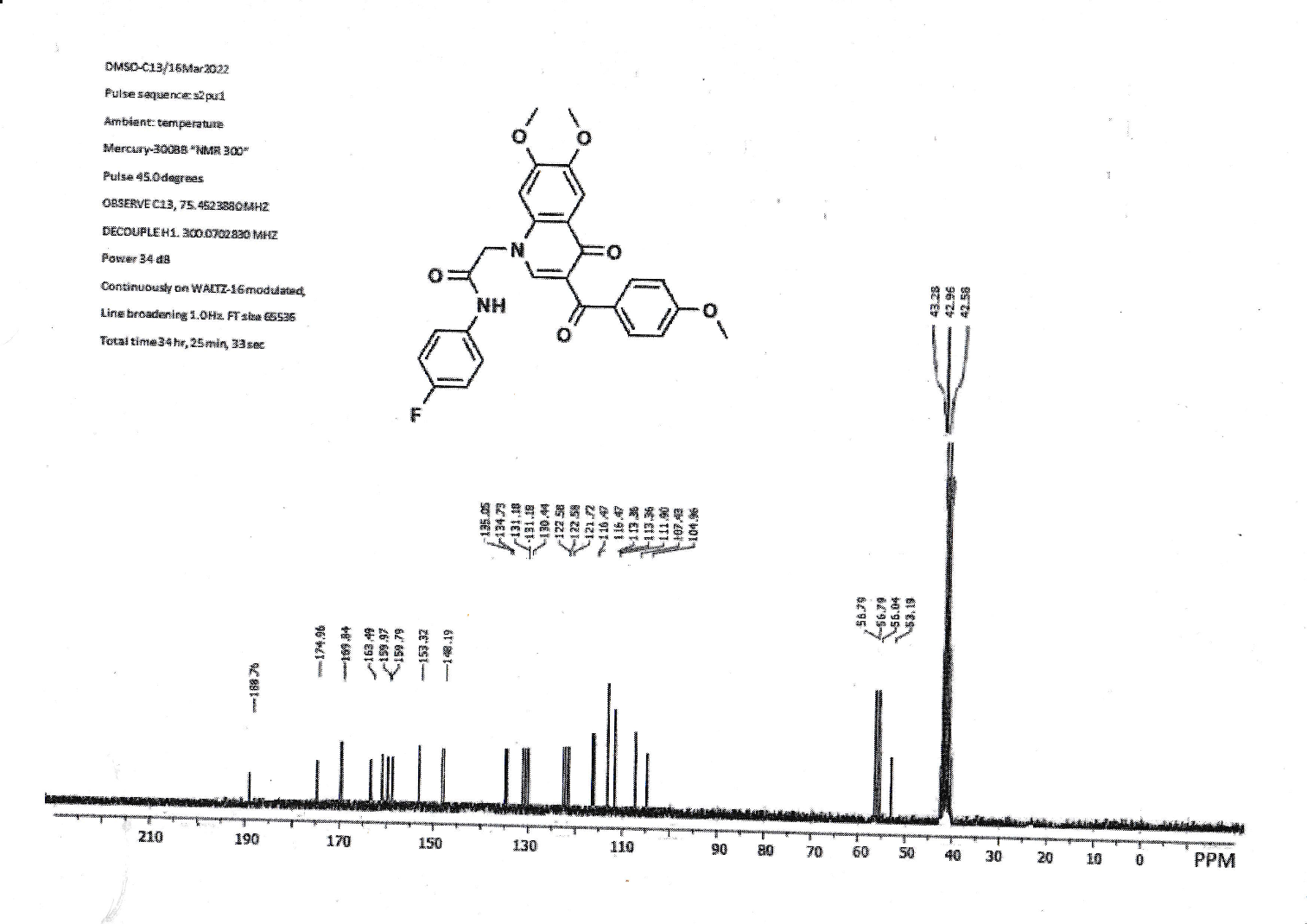

**Figure S8** ^1^H and ^13^C NMR spectrum of compound **8**

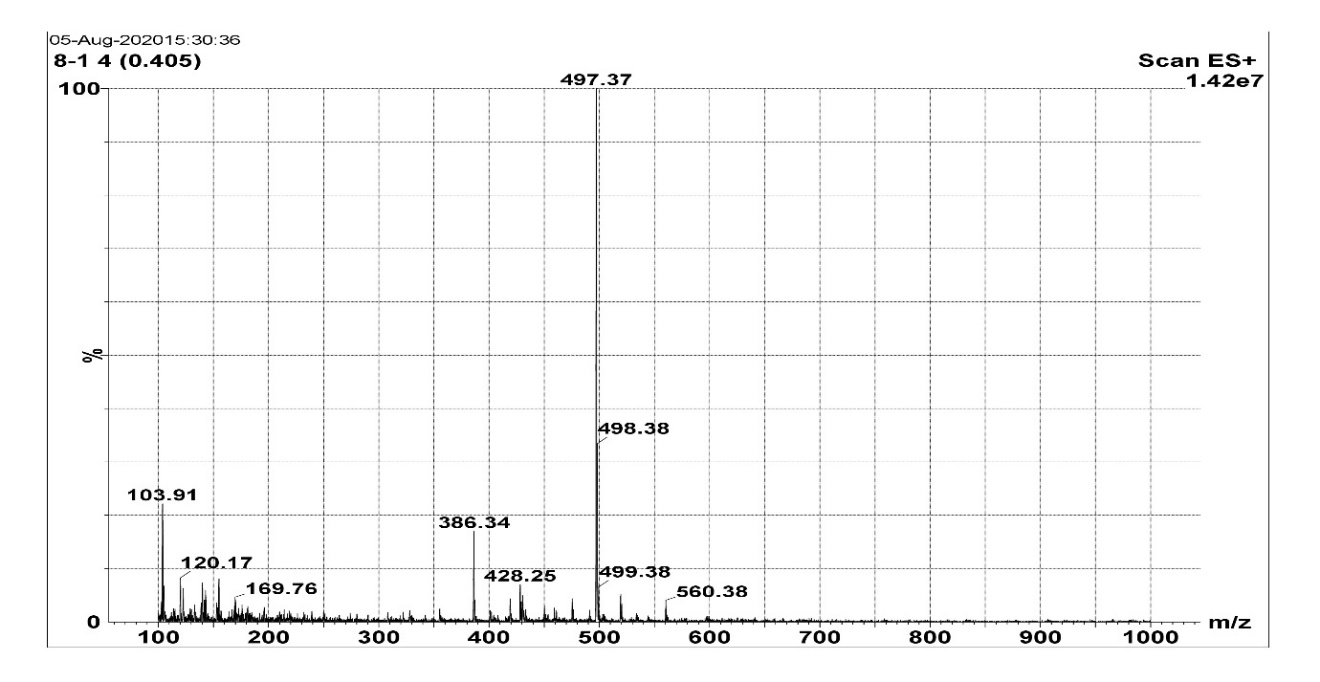

**Figure S9** Mass spectrum of compound **8.**

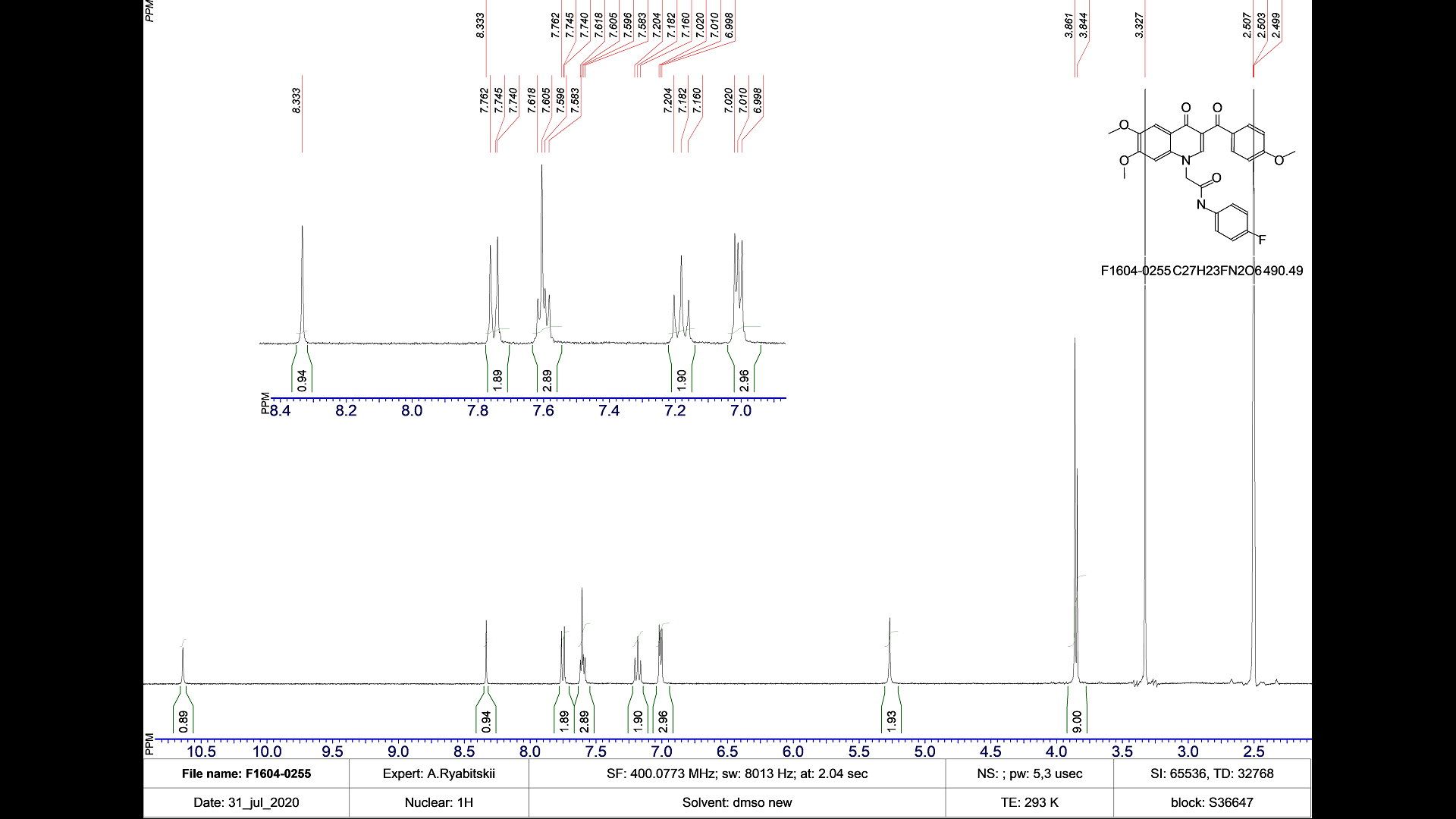

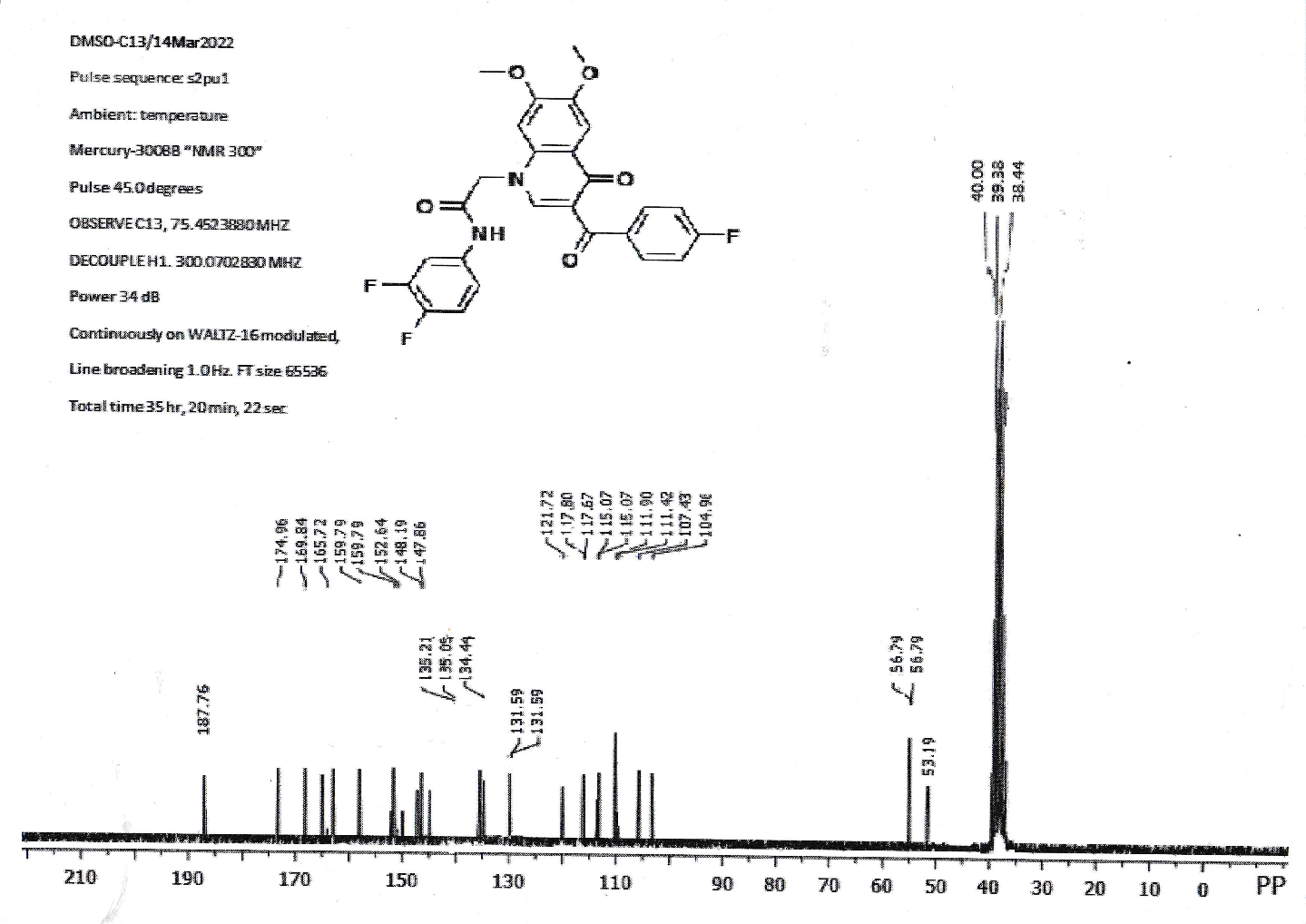

**Figure S10** ^1^H and ^13^C NMR spectrum of compound **9**

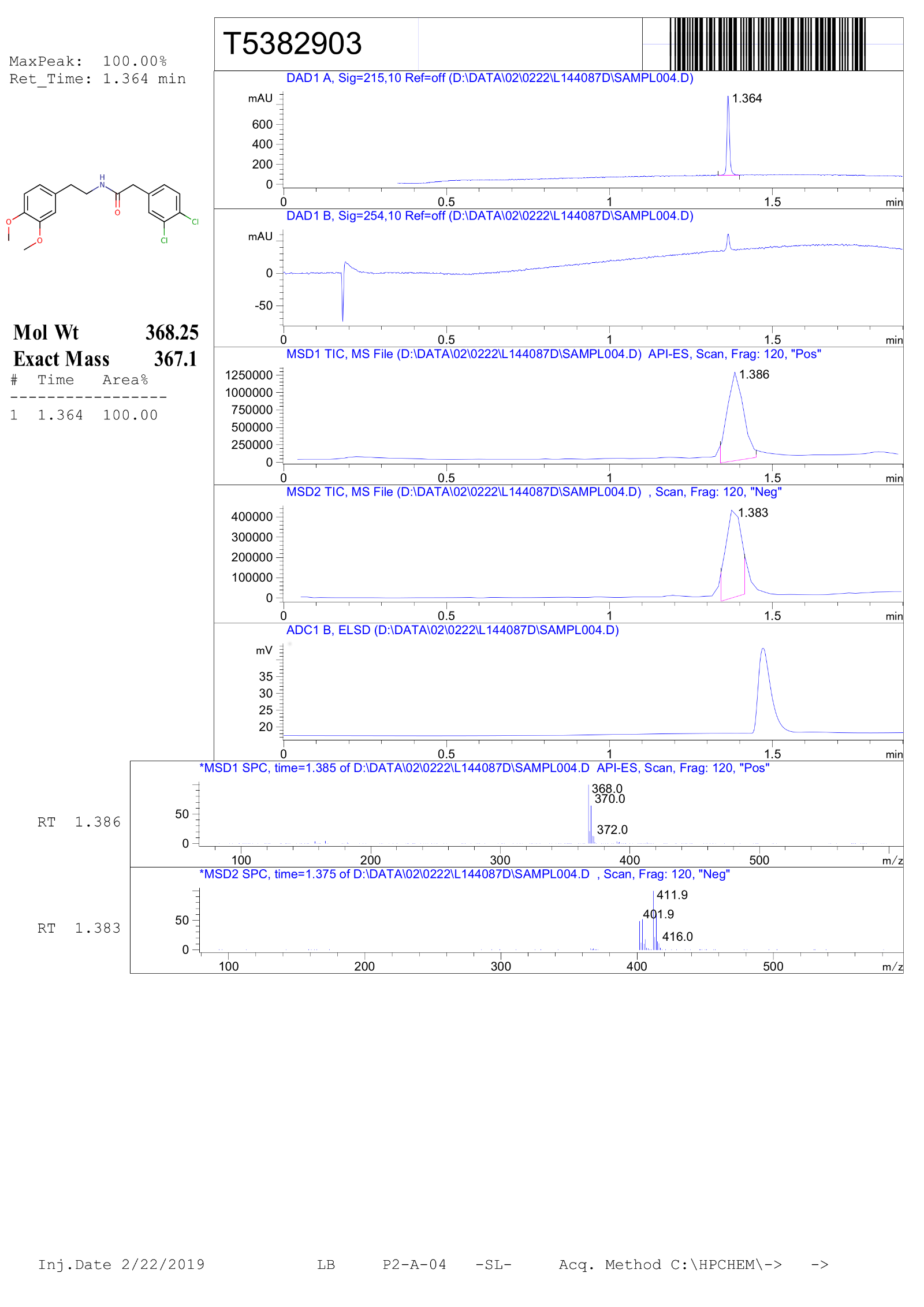

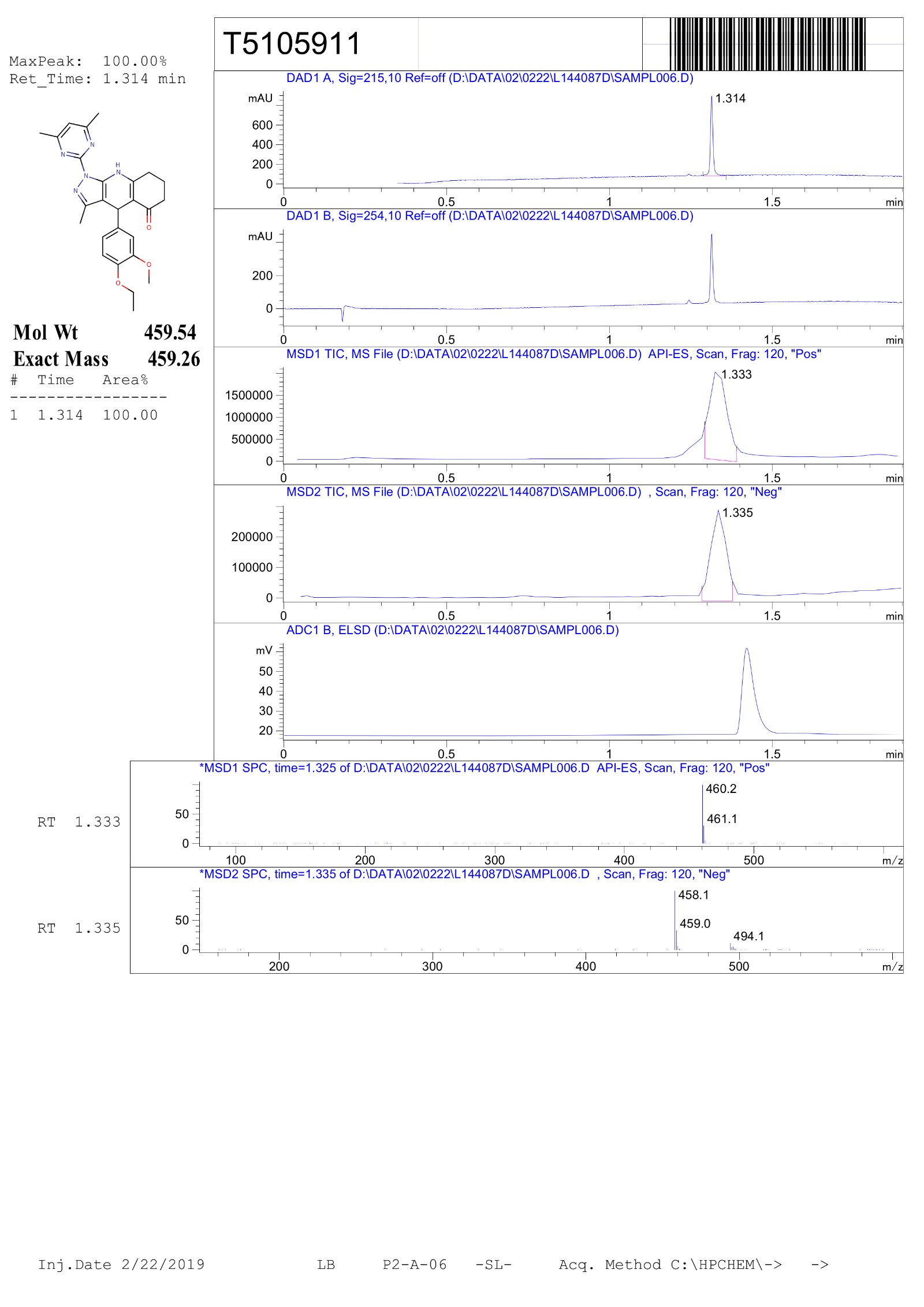

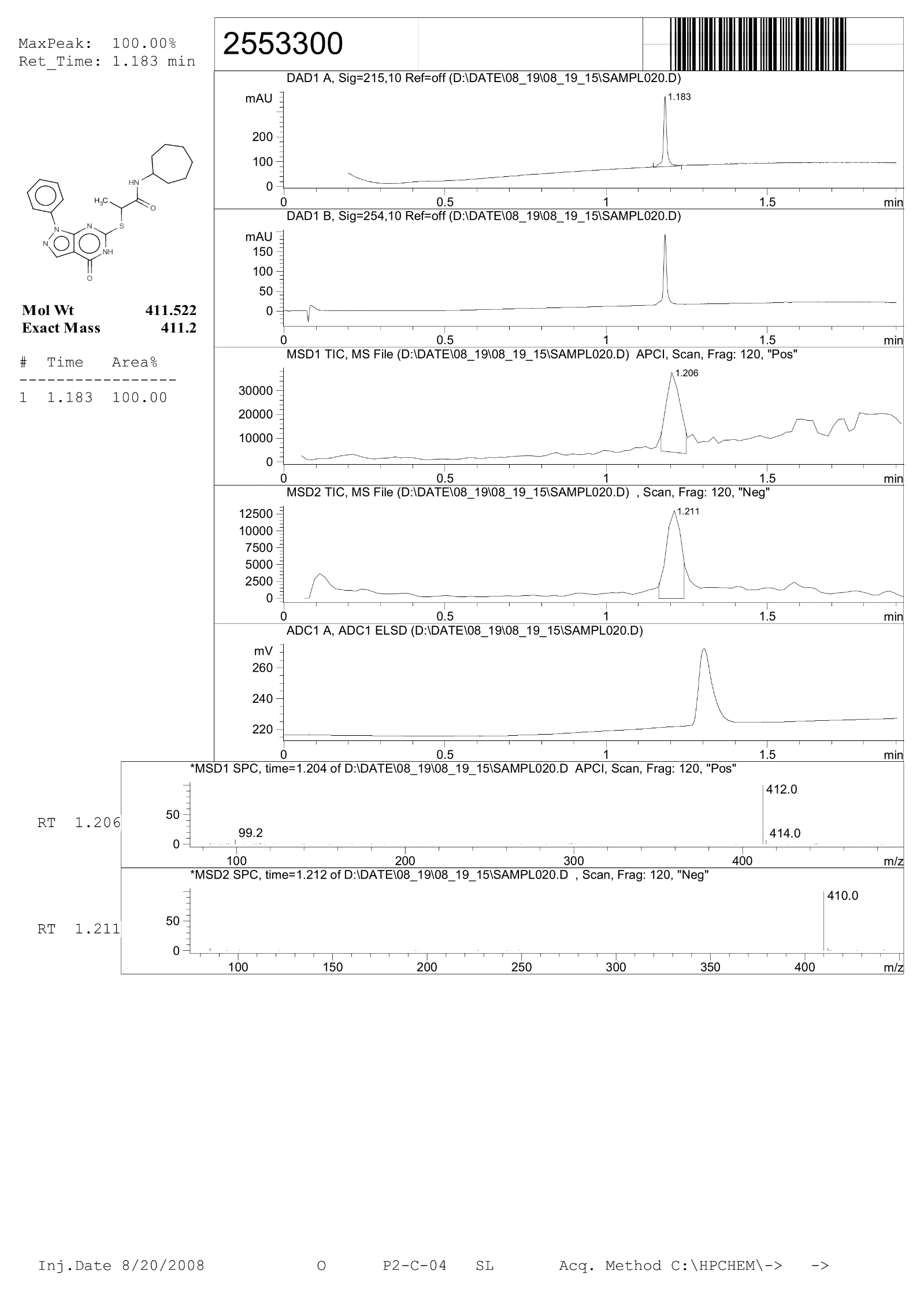

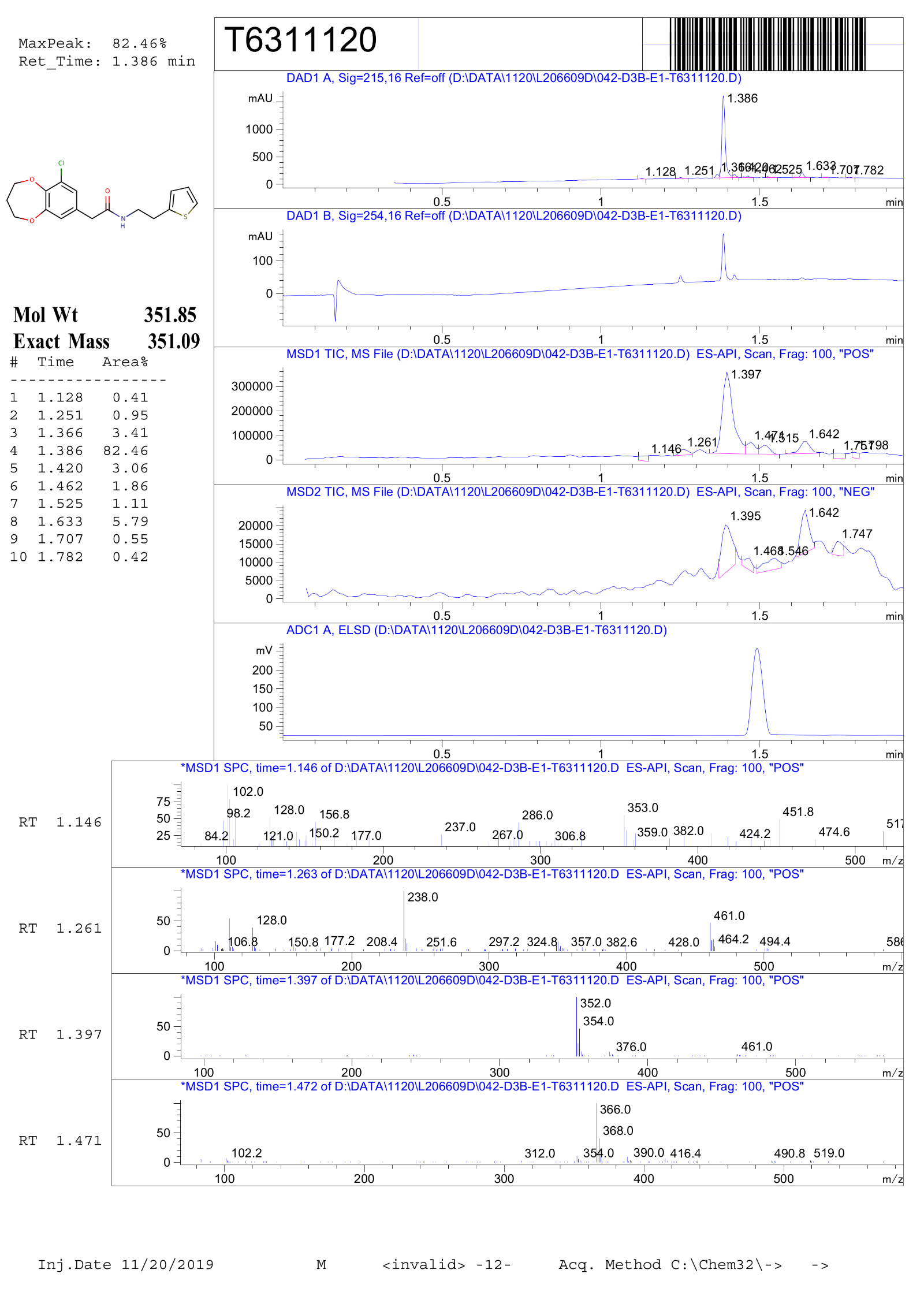

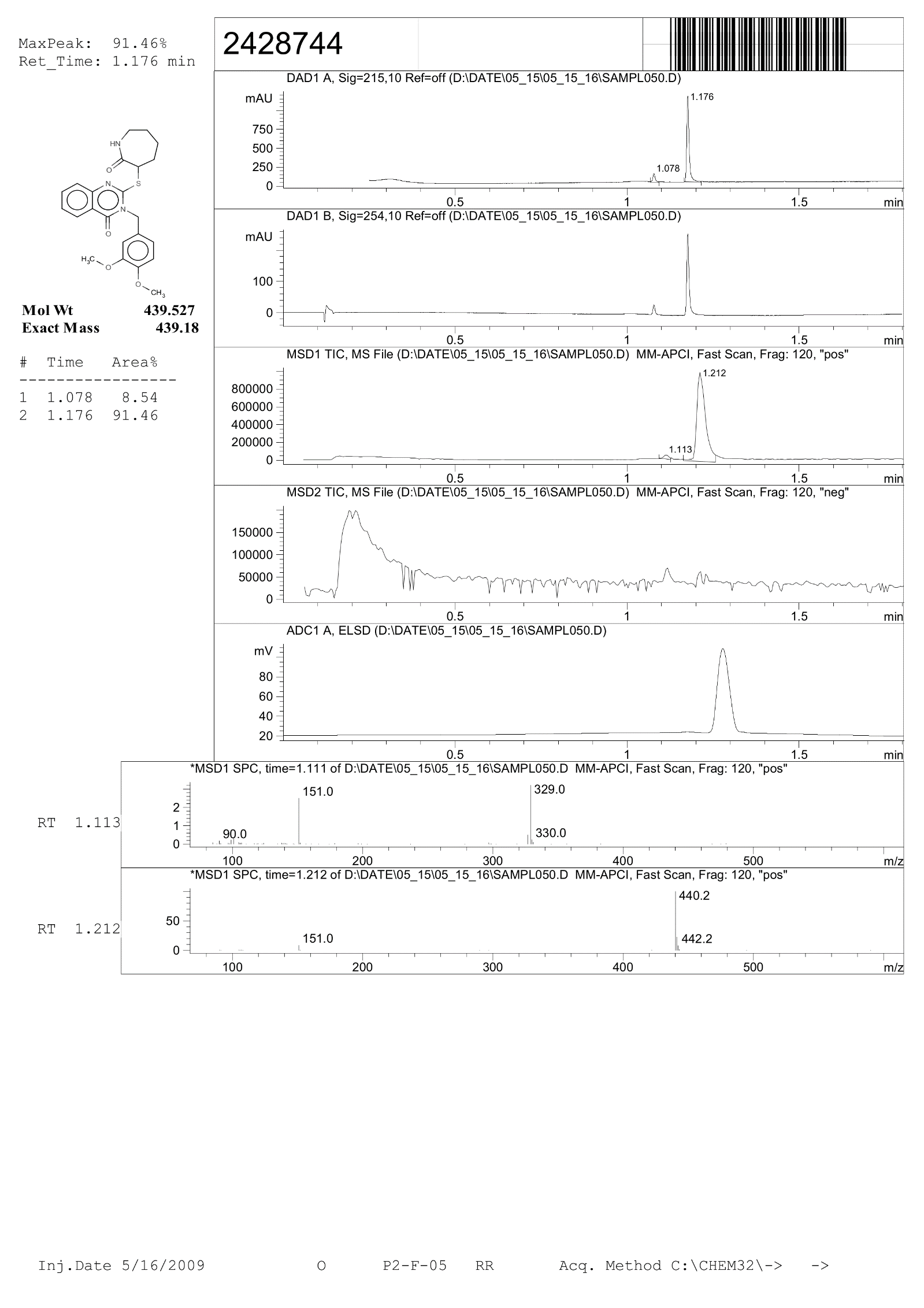

**Figure S11** Spectra and analytical date of the short-listed compounds
